# The Tomato Isochorismate Synthase Coordinates both Salicylic Acid-Mediated Immunity and Phylloquinone-Dependent Chloroplast Function

**DOI:** 10.64898/2026.11.23.768541

**Authors:** Julia Pérez-Pérez, Maria Sulli, Borja Belda-Palazón, Marta Rodríguez-Rodríguez, Francisco Vera-Sirera, Gianfranco Diretto, Ismael Rodrigo, María Pilar López-Gresa, Purificación Lisón

## Abstract

Isochorismate synthase (ICS) provides isochorismate for salicylic acid (SA) and phylloquinone biosynthesis, yet how these functions are integrated in crop species remains poorly understood. We investigated the biological role of the single *ICS* gene in tomato.

CRISPR/Cas9-generated *ics* mutants were analysed using pathogen assays, targeted and untargeted metabolomics, transcriptomics, chloroplast imaging, photosynthetic measurements and polysome profiling.

Loss of ICS strongly reduced pathogen-induced SA accumulation and compromised *Pto*-mediated resistance to *Pseudomonas syringae*. Mutants also displayed severe growth defects, near-complete phylloquinone depletion, impaired photosynthetic performance and profound chloroplast ultrastructural alterations. Supplementation with the phylloquinone precursor 1,4-dihydroxy-2-naphthoic acid partially restored photosynthetic performance. ICS disruption further caused extensive metabolic and transcriptional reprogramming and reduced polysome loading.

These results reveal that the single tomato *ICS* gene contributes to two major outputs of isochorismate metabolism: pathogen-associated SA accumulation and phylloquinone-dependent chloroplast function. Together with emerging evidence for phenylalanine-derived SA biosynthesis in tomato, our findings indicate that SA biosynthetic routes are context-dependent and highlight substantial functional diversification of ICS across plant species.

## Introduction

Plants have evolved a sophisticated immune system based on two interconnected layers of defence. Pattern-triggered immunity (PTI) is activated by the recognition of conserved pathogen-associated molecular patterns, whereas effector-triggered immunity (ETI) involves the recognition of pathogen effectors and generally induces a stronger defence response (Jones and Dangl, 2006). PTI and ETI mutually potentiate each other (Ngou et al., 2021; Yuan et al., 2021) and involve extensive transcriptional, metabolic and hormonal reprogramming, in which salicylic acid (SA) plays a central role (Klessig et al., 2018; Spoel and Dong, 2024).

SA is a major phytohormone regulating immunity against biotrophic and hemi biotrophic pathogens (Loake and Grant, 2007). Upon infection, SA accumulates locally, activating defence-related genes, including PATHOGENESIS-RELATED (PR) genes, promoting hypersensitive cell death and restricting pathogen proliferation (Cameron et al., 1999). SA also plays a central role in systemic acquired resistance (SAR), a long-lasting immune response that protects distal tissues against subsequent infections (Gao et al., 2015).

SA biosynthesis has traditionally been described as occurring through two independent metabolic routes originating from chorismate: the phenylalanine ammonia-lyase (PAL) pathway (El Azaz et al., 2023, Zhu et al., 2025) and the isochorismate (ICS) pathway (Wildermuth et al., 2001; Rekhter et al., 2019, Torrens-Spence et al., 2019). Although both pathways contribute to SA biosynthesis, their relative importance varies among plant species and biological contexts. In *Arabidopsis thaliana*, the isochorismate synthase (*ICS1/SID2*) pathway is the major source of pathogen-induced SA. ICS1 catalyses the conversion of chorismate into isochorismate in chloroplasts, and its disruption severely reduces SA accumulation, compromises defence gene activation and systemic acquired resistance, and increases pathogen susceptibility (Wildermuth et al., 2001; Garcion et al., 2008). However, recent comparative and biochemical evidence has revealed considerably greater diversity in plant SA biosynthesis than previously appreciated. While the *Arabidopsis thaliana* paradigm relies predominantly on the isochorismate pathway, the relative contribution of the ICS-and phenylalanine-derived pathways varies substantially among plant lineages (Ullah et al., 2023; Liu et al., 2026). Notably, two independent studies have recently elucidated the complete phenylalanine-derived SA biosynthetic pathway in rice, providing genetic and biochemical evidence for a major role of this route in other plant species (Zhu et al., 2025; Wang et al., 2025).

Despite these advances in our understanding of SA biosynthesis, considerably less is known about ICS function in crop species (Guo et al., 2023; Wang et al., 2024), including tomato (*Solanum lycopersicum*). Unlike *Arabidopsis thaliana*, which contains two functional ICS genes, the tomato genome contains a single *ICS* gene. Although previous studies using virus-induced gene silencing (VIGS) suggested that tomato ICS contributes to SA accumulation and resistance against *Pseudomonas syringae* pv. *tomato* (Pst), stable loss-of-function mutants have not been available, limiting a comprehensive characterization of ICS biological functions throughout plant development (Uppalapati et al., 2007; Yang et al. 2025). Furthermore, whether tomato primarily relies on the ICS pathway for SA biosynthesis during immunity, as occurs in *Arabidopsis thaliana*, or other alternative pathways such as PAL make substantial contributions, remains unresolved.

Tomato represents an excellent model to investigate SA-dependent immunity because of its well-characterized interaction with Pst (Lonjon et al., 2024). In particular, the resistant cultivar Rio Grande-PtoR, carrying the Pto resistance gene, specifically recognizes the bacterial effector AvrPto, and activates effector-triggered immunity (ETI), providing a well-established genetic system to investigate resistance responses to Pst (Willis and Kinscherf, 2009; López-Gresa et al., 2017).

Beyond its role in salicylic acid biosynthesis, isochorismate also serves as a precursor for phylloquinone (vitamin K₁), an essential component of the photosynthetic electron transport chain. Previous studies in *Arabidopsis thaliana* and crop species have linked ICS activity to phylloquinone biosynthesis, chloroplast development and photosynthetic performance, revealing substantial functional diversification of ICS among plant species (Gawroński et al., 2013; Guo et al., 2023; Wang et al., 2024). In tomato, which contains a single *ICS* gene, whether the same enzyme simultaneously supports SA-dependent immunity and phylloquinone-dependent chloroplast function is still unknown.

Here, we generated the first stable CRISPR/Cas9 knockout mutants of ICS in tomato and performed an integrated physiological, biochemical, transcriptomic, metabolomic and polysome-profiling characterization to investigate its contribution to immunity and cellular homeostasis. Beyond confirming the critical role of ICS in SA accumulation during ETI and resistance to Pst, our analyses reveal distinct alterations in chloroplast development, photosynthetic metabolism, lipid homeostasis and volatile biosynthesis, together with previously unexplored changes in translational regulation. Together, our results identify tomato ICS as a central metabolic hub linking chloroplast function, growth and immune signalling, providing an integrated view of its biological functions beyond salicylic acid biosynthesis in tomato.

## Results

### The tomato Isochorismate Synthase Is Required for Full Pathogen-Induced Salicylic Acid Accumulation and Pto-mediated Resistance to *Pseudomonas syringae*

To study the role of ICS and SA in defence, we generated CRISPR/Cas9 ICS knockout mutants in tomato cv. ‘Rio Grande’ to investigate the contribution of ICS in SA biosynthesis and plant immunity during virulent and avirulent infections with Pst.

The tomato genome contains a single ICS gene (Solyc06g071030.2), which was targeted using four CRISPR/Cas9 guide RNAs. Two independent mutant lines were obtained, carrying deletions of 130 bp (ics1) and 195 bp (ics2) (Figures 1A and S1). Sequence and cDNA analyses revealed altered coding sequences resulting in predicted truncated proteins of 535 and 514 amino acids, respectively, compared with 580 amino acids in WT. Both mutant lines exhibited severe developmental defects and could not be maintained under greenhouse conditions, requiring continuous in vitro culture with sucrose supplementation, consistent with previous observations (Yang et al., 2025; Figure 1A).

**Figure 1.**
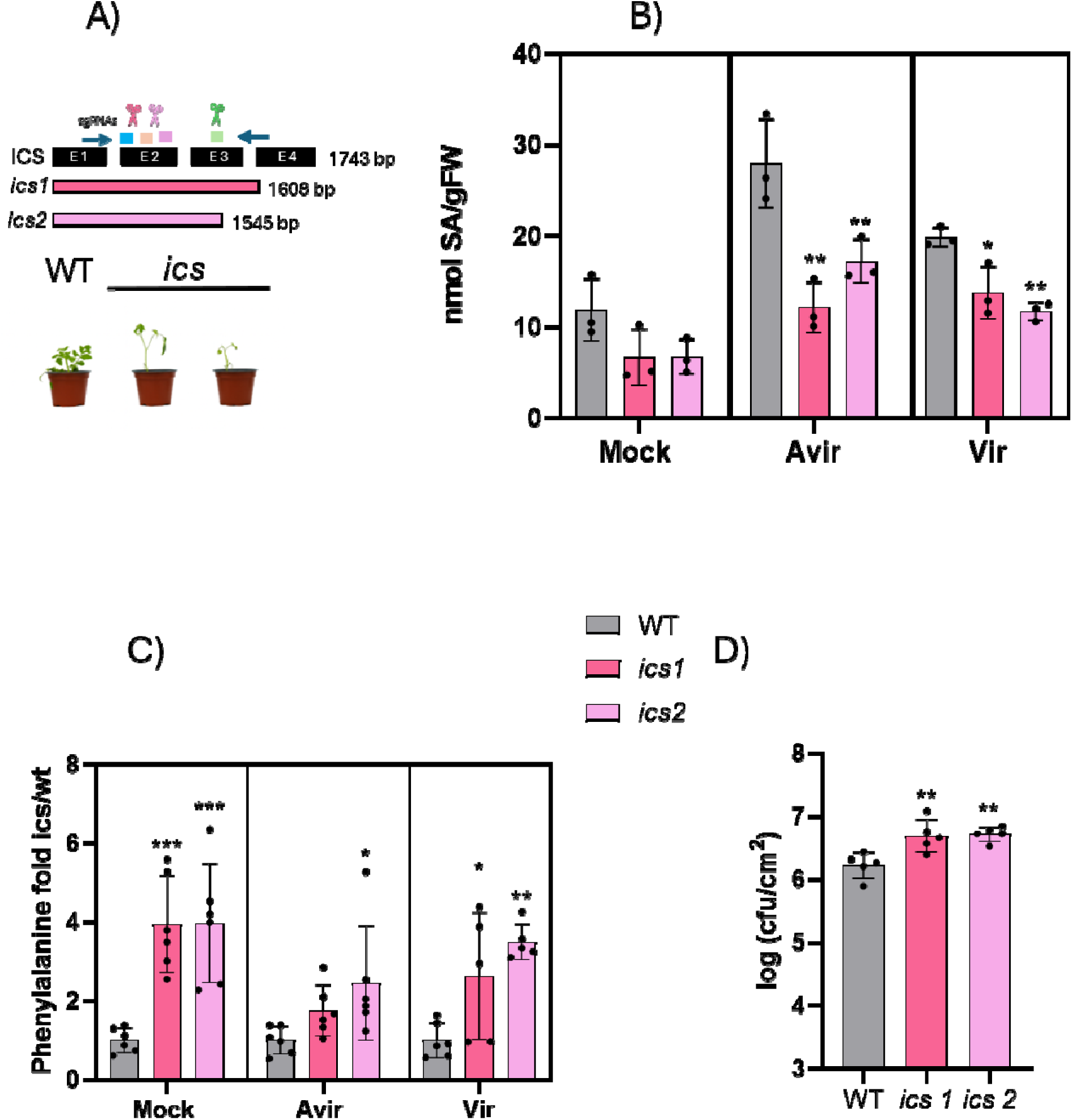
Generation and characterization of CRISPR/Cas9 tomato *ics* mutant lines. **(A)** Schematic representation of the tomato *ICS* gene structure showing four exons (E1-E4) and the positions of the three designed single-guide RNAs (sgRNAs). Two independent deletions generated by sgRNA1, sgRNA2 and sgRNA3 are indicated, resulting in the *ics1* (130 bp deletion) and *ics2* (195 bp deletion) mutant alleles. Phenotype of wild-type (WT) and *ics* mutant plants grown under greenhouse conditions is shown below. **(B)** Endogenous salicylic acid (SA) levels under mock conditions and following virulent or avirulent *Pseudomonas syringae* pv. tomato (Pst) infection (24 hpi). **(C)** Phenylalanine accumulation in *ics* plants relative to WT plants, with WT levels normalized to 1, mock conditions and following virulent or avirulent Pst infection **(D)** Bacterial growth of Pst during avirulent interactions. Data represent the mean ± SD of biological replicates (n=5 for Pst growth, n=3 for metabolites). Asterisks indicate statistically significant differences compared with WT (*p* < 0.05, \**p* < 0.01, \*\**p* < 0.001; one-way ANOVA followed by Dunnett’s multiple comparison test).

To test the contribution of ICS to SA biosynthesis in tomato, total SA levels were quantified under basal and biotic stress conditions (Figure 1B). Constitutive SA levels in the mock mutants tended to be lower than in wild type plants, although this reduction was not statistically significant. Under biotic stress, SA accumulation was analysed following infection with two strains of Pst: a virulent strain (leading to disease) and an avirulent strain (leading to ETI). During both infections, SA accumulation was markedly impaired in the *ics* mutants, whereas wild-type plants showed a strong induction of SA. These results suggest that *ICS* plays a critical role in SA biosynthesis during virulent and avirulent tomato-Pst interactions.

Since SA can also be synthesized via the phenylalanine ammonia-lyase (PAL) pathway, the levels of phenylalanine were also measured (Figure 1C). Interestingly, under basal conditions and during virulent interactions, phenylalanine levels were elevated in the mutants, suggesting that loss of *ICS* alters the pool of this aromatic amino acid. In contrast, this increase was not observed during avirulent interactions, when SA accumulation was almost completely abolished in the mutants, highlighting the essential contribution of ICS to SA biosynthesis during ETI.

Finally, we assessed whether disruption of ICS and the associated reduction in SA during avirulent interactions were accompanied by impaired disease resistance (Figure 1D). Both *ics* mutant lines displayed significantly increased bacterial growth compared with wild-type plants following infection with the avirulent Pst strain. The enhanced bacterial proliferation was accompanied by a clear loss of the resistance phenotype associated with the Pto-mediated interaction, indicating that ICS is required for full resistance to avirulent Pst.

Together, these findings establish *ICS* as a major contributor to pathogen-induced SA accumulation in tomato and demonstrate that its disruption compromises Pto-mediated resistance.

### Loss of *ICS* Triggers Broad Constitutive Reprogramming of Immune and Cellular Homeostasis Pathways

Given the severe developmental defects and growth arrest displayed by the *ics* mutant lines, we investigated the molecular consequences of *ICS* disruption through transcriptome profiling under basal, non-infected conditions. RNA-seq analysis revealed profound changes in gene expression, indicating that loss of *ICS* affects cellular functions (Figure 2).

**Figure 2.**
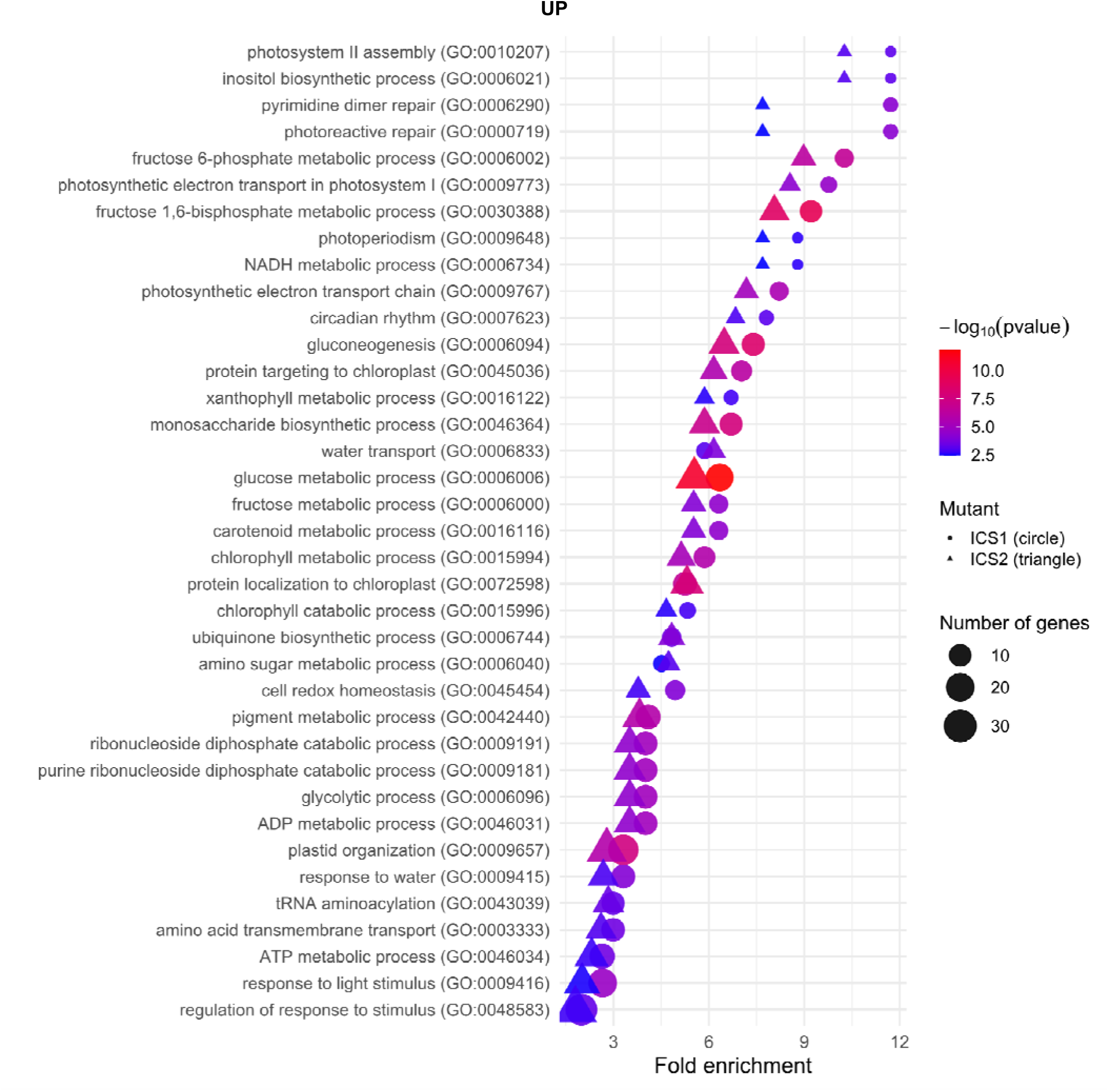
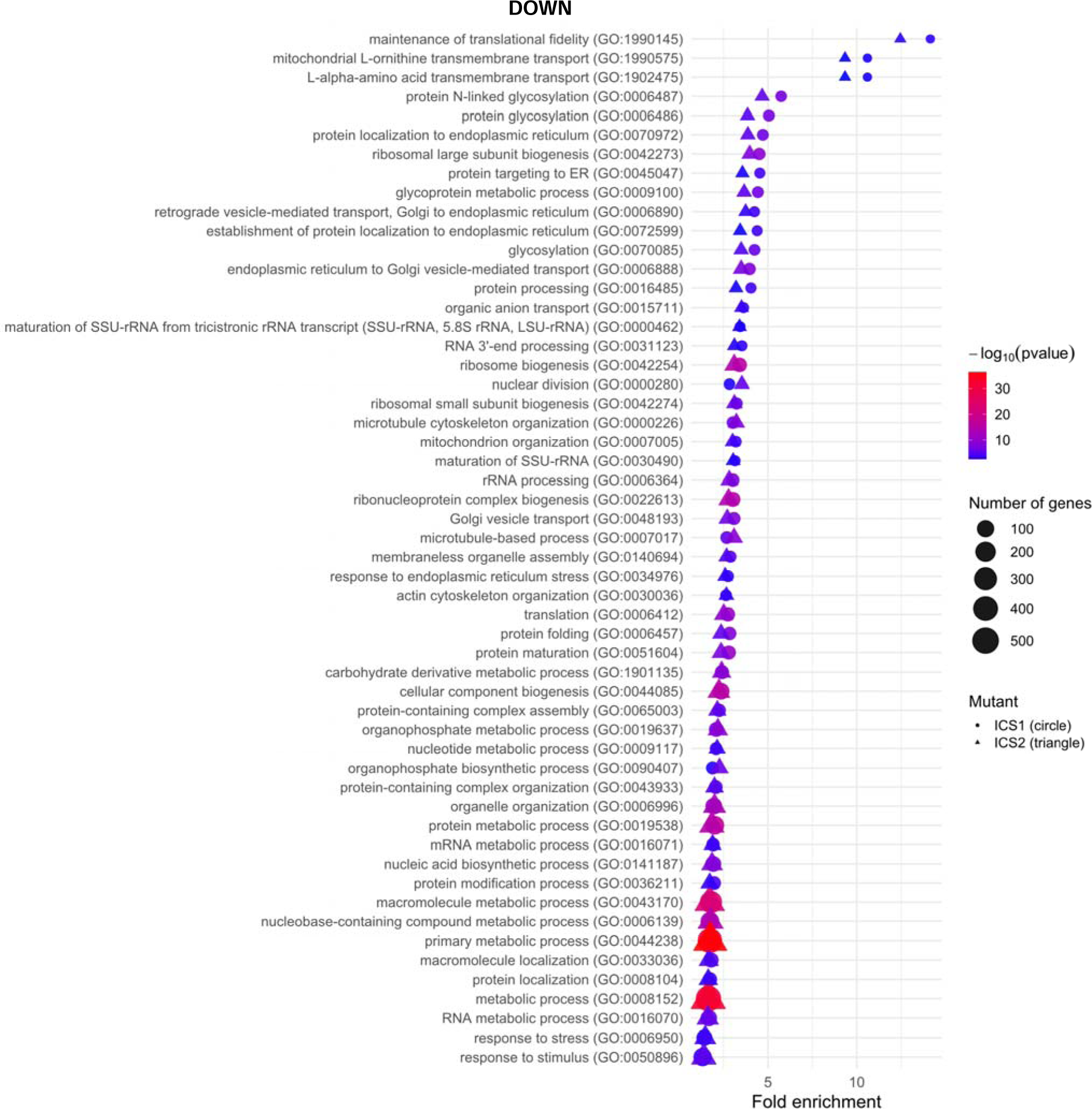
Transcriptomic reprogramming in CRISPR-generated *ics* mutant tomato plants under basal conditions. UP **(A)** and DOWN **(B)** Gene Ontology (GO) enrichment analysis of differentially expressed genes shared by both mutant lines identified significant enrichment of biological processes related to primary metabolism. Bubble plot showing significantly enriched GO biological process terms among the differentially expressed genes shared by both *ics* mutant lines relative to wild-type plants. The x-axis represents fold enrichment of each GO term, while bubble size corresponds to the number of genes associated with each category. Bubble color indicates statistical significance (−log10 *p*-value). Circles represent *ics1* and triangles represent *ics2* mutants.

Principal component analysis (PCA) revealed a clear separation between wild-type and *ics* mutants, with PC1 explaining 60% of the variance (Figure S2A). Differential expression analysis confirmed extensive transcriptional reprogramming in both mutant lines (Figure S2B). Notably, *PR1* and *P23* were significantly downregulated in both mutants, as shown by RNA-seq and validated by RT-qPCR (Figure S2C).

To further characterize the global transcriptional changes associated with ICS disruption, Gene Ontology (GO) enrichment analysis was performed on differentially expressed genes. Upregulated genes in the *ics* mutants were enriched in processes associated with primary metabolism, energy production, and stress adaptation. Notably, terms related to photosynthesis, including photosystem I and II assembly, photosynthetic electron transport, and light reactions, were significantly overrepresented. This was accompanied by the induction of multiple metabolic pathways, such as carbohydrate metabolism (e.g. glucose metabolic process, gluconeogenesis; Figure S3), amino acid catabolism, and lipid oxidation, suggesting a metabolic reprogramming toward energy mobilization (Figure 2A).

In contrast, analysis of downregulated genes revealed a widespread enrichment of terms associated with core cellular functions (Figure 2B), including translation, ribosome biogenesis, RNA processing, and gene expression, indicating a general reduction in biosynthetic and growth-related activities. Processes linked to cytoskeleton organization (e.g. actin filament organization and microtubule-based processes), cell cycle progression, and intracellular transport were also significantly affected. Together, these results suggest that *ICS* loss leads to a broad alteration of cellular homeostasis and growth-associated functions, in addition to compromising immune signalling.

Overall, the GO enrichment analysis reveals that *ICS* disruption alters the transcriptional landscape of tomato plants, affecting both cellular processes and metabolic functions. The downregulation of genes involved in protein synthesis, RNA processing, cytoskeleton organization, cell cycle progression, and intracellular transport points to a reduction in growth-associated and biosynthetic activities. In parallel, the induction of photosynthetic, carbohydrate, amino acid, and lipid metabolic processes suggests a substantial reorganization of energy and metabolic pathways. Thus, rather than producing a specific alteration in SA-related defence responses, loss of ICS is associated with a broad transcriptional reprogramming that extends across fundamental cellular and metabolic processes, highlighting the central role of ICS in maintaining the balance between defence, growth, and cellular homeostasis.

### *ICS* is Essential for Phylloquinone Biosynthesis and Photosystem II Functionality

Given the enrichment of photosynthesis-related genes and the impaired growth of ics mutants, we investigated photosynthetic performance. PSII quantum efficiency was significantly reduced in both mutant lines (Figure 3A), indicating impaired photosynthetic function.

**Figure 3.**
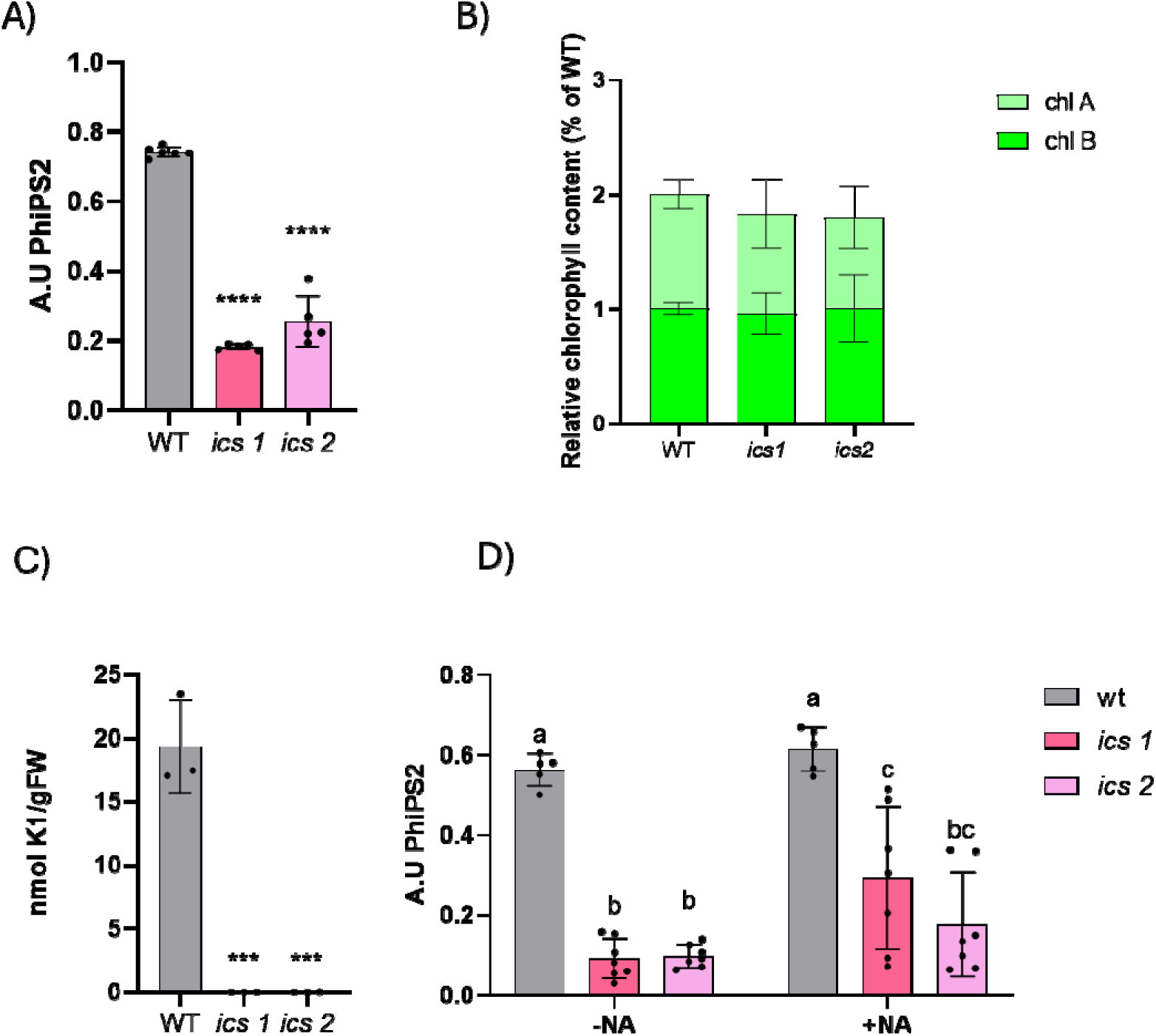
ICS disruption impairs phylloquinone accumulation and photosynthetic performance in tomato. **(A)** Photosystem II (PSII) quantum efficiency in wild-type (WT) and *ics* mutant plants. **(B)** Relative chlorophyll a and b contents determined by HPLC-PDA. **(C)** Phylloquinone (vitamin K1) content quantified by HPLC-fluorescence. **(D)** Analysis of Photosystem II function (arbitrary units, A.U.) before and after a treatment with NA (1,4-dihydroxy-2-naphtoic acid) in wildtype plants and both mutant lines. Data represent mean ± SD. Statistical significance was determined relative to WT using one-way ANOVA followed by Tukey’s multiple comparison test (\**p* < 0.05,** *p* < 0.01,*** *p* < 0.001).

To investigate the biochemical basis of the photosynthetic alterations observed in *ics* mutants, we first quantified chlorophyll levels by HPLC-PDA (Figure 3B). Although mutant plants occasionally exhibited subtle differences in leaf coloration compared with the WT, these changes were not consistently associated with reduced chlorophyll accumulation. Indeed, chlorophyll levels were slightly decreased whereas no significant differences were detected.

We next examined phylloquinone, an essential electron carrier in photosystem I whose biosynthesis depends on isochorismate. HPLC-FL (Figure 3C) and HPLC-MS analyses revealed a drastic reduction in phylloquinone levels in *ics* mutants, reaching almost undetectable levels. This reduction was also maintained following infection with either virulent or avirulent Pst (Figure S4). Together, these results indicate that ICS deficiency severely compromises phylloquinone accumulation without consistently affecting total chlorophyll levels.

To further test whether the impaired photosynthetic performance of the *ics* mutants resulted from a defect in phylloquinone biosynthesis, we performed a metabolic complementation experiment using 1,4-dihydroxy-2-naphthoic acid (NA) (Römer et al., 2024; Wang et al., 2024; Kim et al. 2008), an intermediate in the phylloquinone biosynthetic pathway which is located downstream of the dependent isochorismate step. As Figure 3D indicates, NA supplementation partially restored CS-PSII quantum efficiency in the *ics* mutants compared with untreated mutant plants. This partial recovery indicates that providing a phylloquinone precursor can alleviate the photosynthetic defect caused by *ICS* disruption, supporting a functional link between impaired ICS-dependent phylloquinone biosynthesis and the reduced photosynthetic performance observed in the mutant lines.

Overall, these results show that ICS is required for phylloquinone accumulation and normal photosynthetic function in tomato. The upregulation of photosynthesis-related genes in the *ics* mutants may therefore represent a compensatory transcriptional response to impaired photosynthetic performance.

### *ICS* Disruption Severely Affects Chloroplast-associated Lipid Metabolism and Chloroplast Ultrastructure

To investigate the metabolic basis of the severe developmental phenotype observed in the *ics* mutants, we performed an untargeted analysis of the apolar metabolome. Again, the Principal component analysis (PCA) given by the general metabolic distribution within samples revealed a striking separation between wild-type and mutant plants (Figure 4A), with PC1 explaining approximately 45.6% of the total variance. Both *ics1* and *ics2* clustered separately from the wild type, indicating that disruption of ICS profoundly affects the apolar metabolic fraction

**Figure 4.**
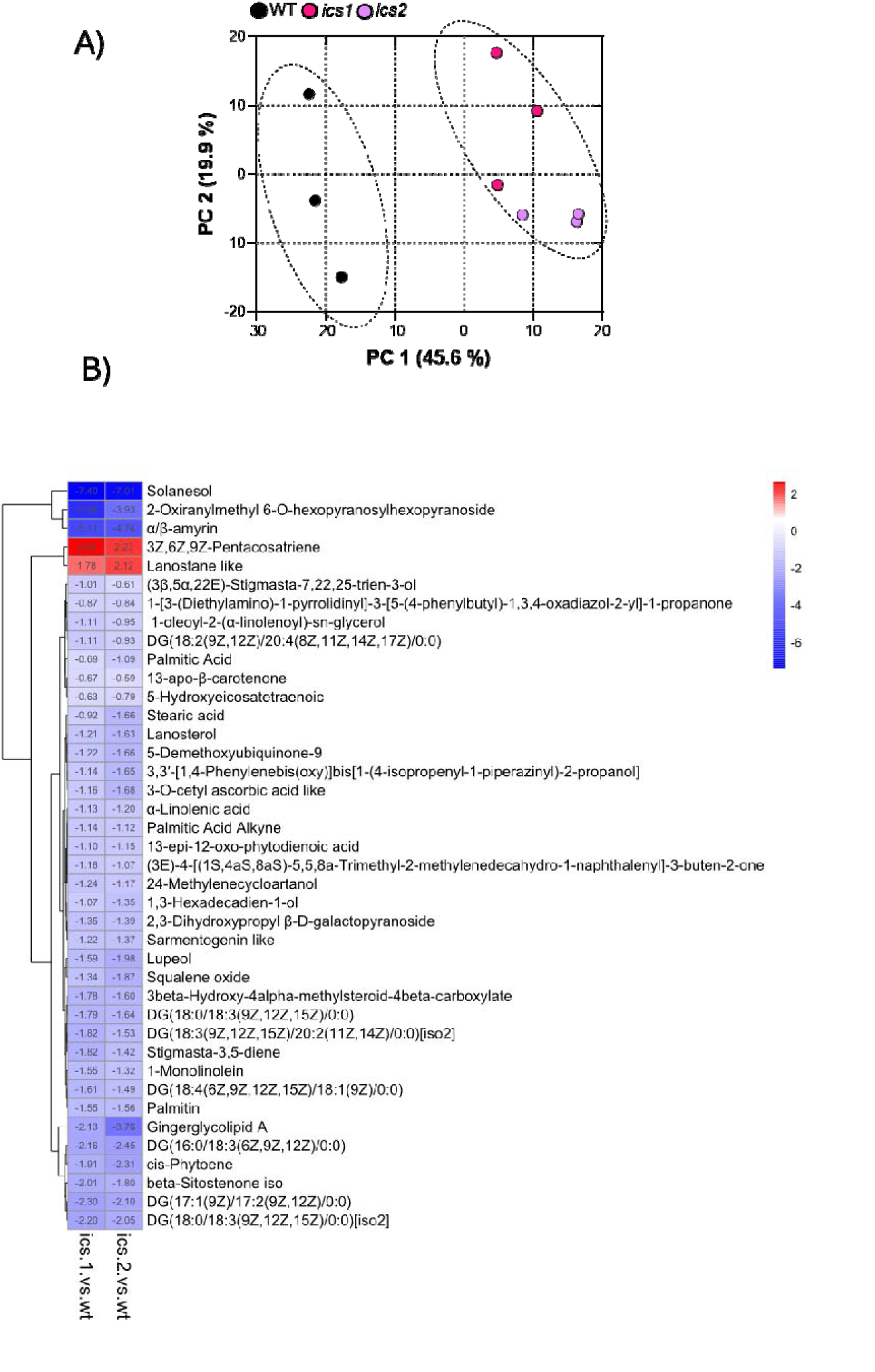
ICS disruption profoundly alters the non-polar metabolome. **(A)** Principal component analysis (PCA) of untargeted LC-MS non-polar metabolite profiles showing a clear separation between wild-type (WT) and both *ics* mutant lines along PC1. **(B)** Hierarchical clustering heatmap of the most significantly altered metabolites. Color intensity represents the fold change relative to WT (*ics*/WT), ranging from dark blue for strongly down-regulated/depleted metabolites to red for highly over-accumulated ones.

Untargeted non-polar metabolomics revealed a clear separation between WT and *ics* mutants, with PC1 explaining 45.6% of the variance (Figure 4A). Among 517 detected metabolic features, 52 were significantly altered in both mutants, of which 40 were tentatively identified by accurate mass and MS/MS analysis. Most of compounds showed decreased levels in *ics* mutants compared to the WT (Volcano plots, Figure S5).

Non-polar metabolomic analysis revealed significant alterations in lipid and carotenoid metabolism in *ics* mutants (Figure 4B, Table S1). Several diacylglycerols (DAGs) and monoacylglycerols (MAGs) (Yu et al., 2023), together with fatty acids involved in thylakoid membrane formation, including α-linolenic, palmitic and stearic acids, were markedly reduced. Carotenoid-related metabolites, including cis-phytoene and 13-apo-β-carotenone, also decreased (Figure 4B). Given the structural and photoprotective roles of these compounds in chloroplasts (Havaux, 1998), their coordinated reduction is consistent with impaired thylakoid membrane organization and photosynthetic function.

To determine whether these metabolic defects were accompanied by structural alterations, chloroplast organization was examined by confocal microscopy based on chlorophyll autofluorescence (Figures 5A and B). Wild-type plants displayed predominantly compact chlorophyll fluorescence signals, whereas both *ics1* and *ics2* mutants exhibited a higher prevalence of ring-like fluorescence patterns with less fluorescent central regions.

**Figure 5.**
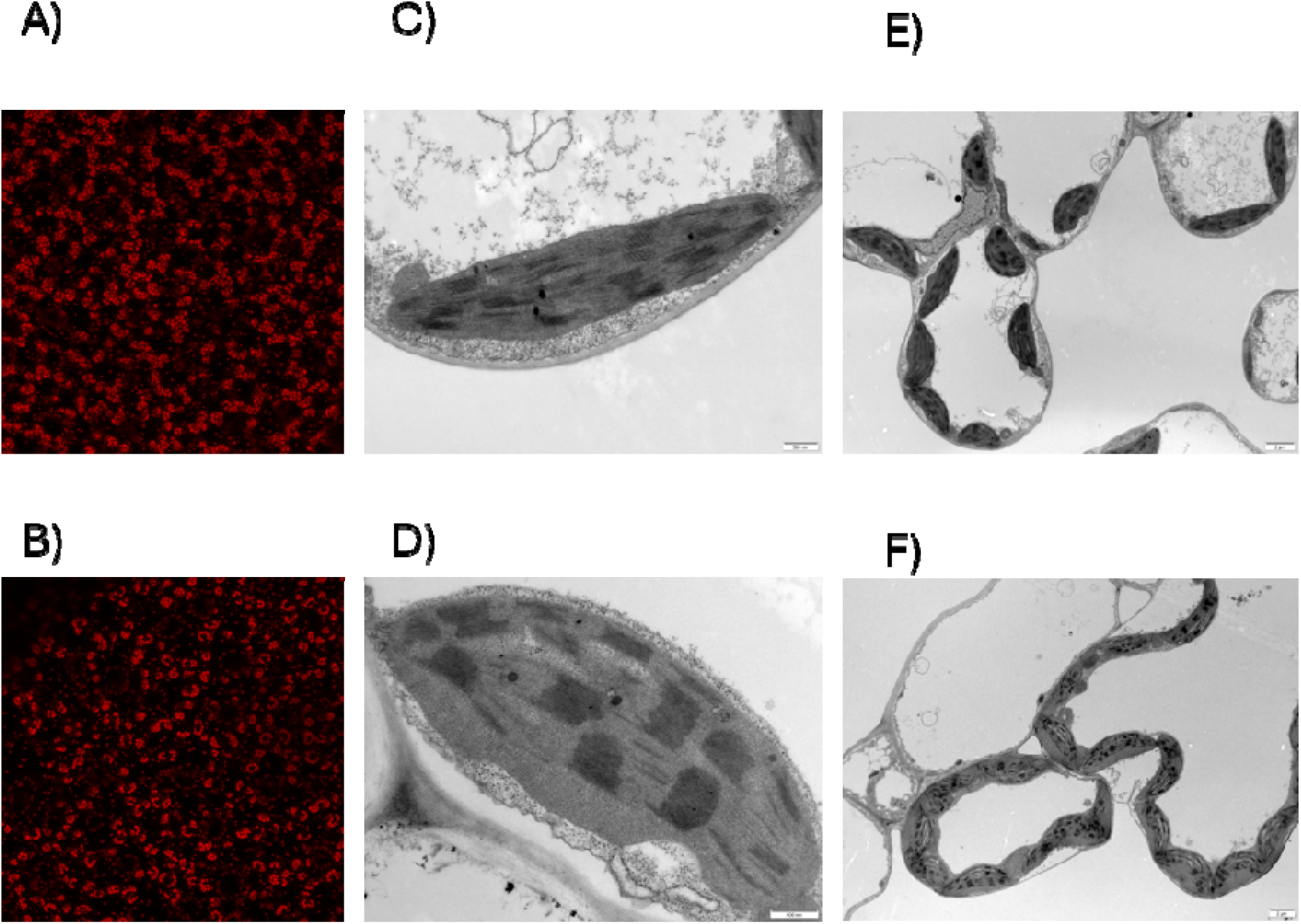
Analysis of chloroplast reveal changes between *ics* and wt plants. (A, B) Confocal laser scanning microscopy of chlorophyll autofluorescence in WT and both mutants *ics* lines (C, D, E, F) Ultrastructural analysis of chloroplasts in wild-type and *ics* mutant plants by transmission electron microscopy (TEM). Representative TEM micrographs of leaf mesophyll cells and chloroplasts from *ics* mutants (upper panels) and wild-type (WT) plants (lower panels). Low-magnification images show the overall cellular organization, while high-magnification images highlight chloroplast ultrastructure. Representative images from WT and the *ics1* mutant are shown for both techniques (the same phenotype was also observed in the *ics2* mutant). Scale bars are indicated in each panel.

To further characterize these alterations at higher resolution, chloroplast ultrastructure was analysed by transmission electron microscopy (TEM) (Figures 5C-F). Wild-type chloroplasts displayed the characteristic lenticular morphology, with tightly stacked grana and an organized thylakoid network. In contrast, chloroplasts from both *ics* mutants exhibited a more rounded and swollen morphology, accompanied by partial disorganization of the thylakoid membranes.

Taken together, these results demonstrate that loss of ICS function extends far beyond SA biosynthesis and immune signalling. ICS disruption causes a severe depletion of chloroplast-associated lipids, carotenoids, prenyl lipids, and thylakoid fatty acids, ultimately leading to abnormal chloroplast development and organization. These defects likely contribute to the pale green phenotype, impaired growth, and compromised physiological performance observed in the *ics* mutant lines.

### *ICS* Disruption Causes Broad Metabolic and Hormonal Reprogramming in Tomato

To assess the broader metabolic consequences of ICS disruption, we analysed volatile and semi-polar metabolites in WT and ics plants. Both analyses clearly separated mutant and WT samples (Figure 6A,B).

**Figure 6.**
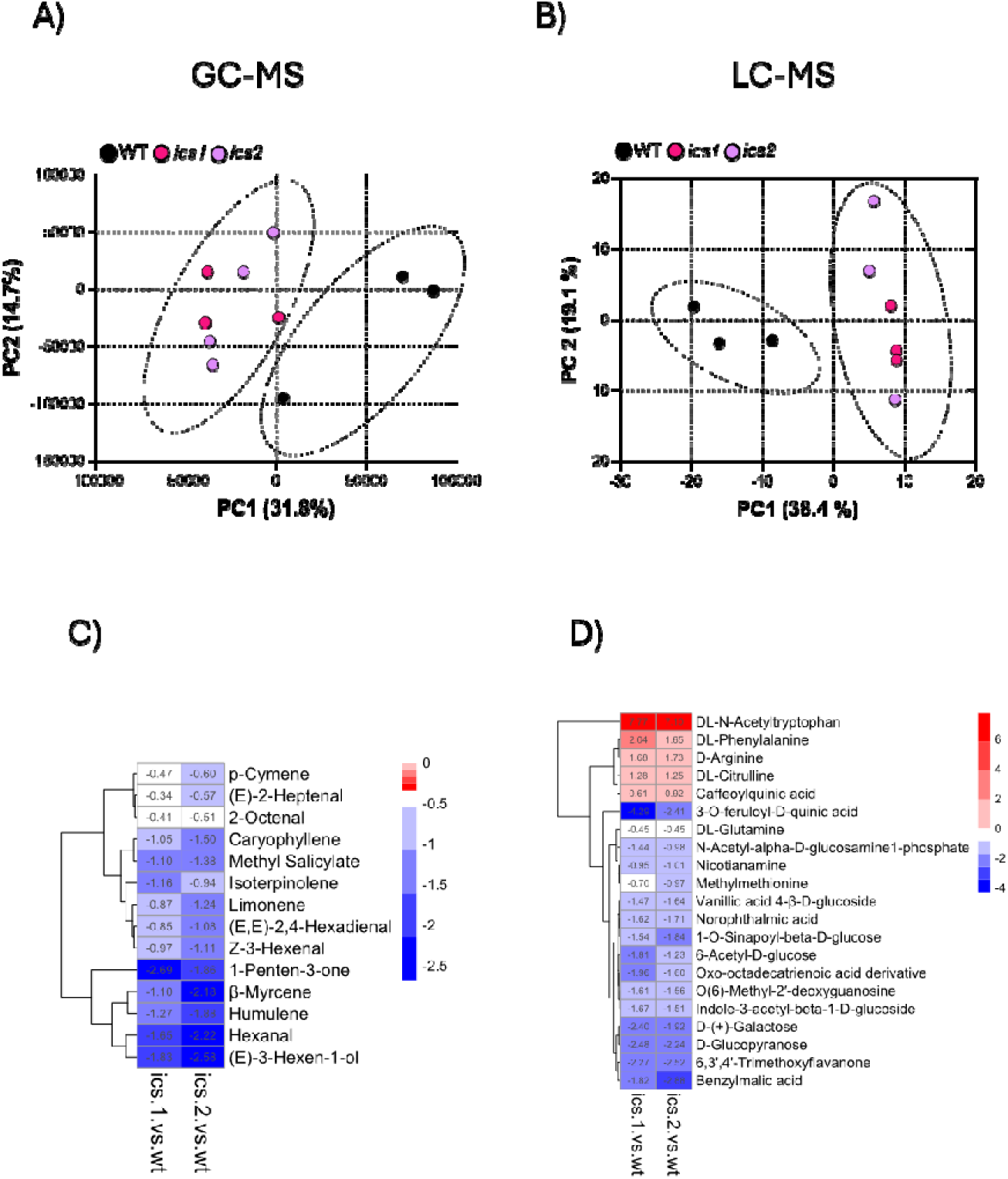
Disruption of *ICS* alters the constitutive volatile and semi polar metabolite profiles of tomato plants. **(A)** Principal component analysis (PCA) of volatile organic compounds (VOCs) detected by GC-MS in wild-type (WT), *ics1*, and *ics2* tomato leaves under mock conditions. **(B)** Principal component analysis (PCA) of untargeted semi-polar metabolites detected by LC-MS in WT, *ics1*, and *ics2* plants under basal conditions. **(C)** Heatmap of the differentially accumulated volatile compounds identified by GC-MS. Columns represent the log_₂_ fold change (*ics1*/WT and *ics2*/WT). Blue indicates reduced accumulation and red indicates increased accumulation relative to WT. **(D)** Heatmap of the most significantly altered semi-polar metabolites identified by untargeted LC-MS. Heatmaps show log_₂_ fold changes (ics1/WT and ics2/WT), with red indicating increased abundance, blue decreased abundance and white no change relative to WT. Values within cells indicate log_₂_ fold changes. Metabolites were hierarchically clustered by row, while columns retained their original order.

Among the affected VOC classes, sesquiterpenes were consistently reduced, including caryophyllene and humulene, whereas the monoterpene limonene was also decreased, together with β-myrcene and isoterpinolene (Figure 6C). These changes suggest that disruption of ICS affects both the mevalonate and plastidial isoprenoid pathways involved in volatile biosynthesis. In addition, several green leaf volatiles (GLVs), including (*E*)-3-hexen-1-ol, (*Z*)-3-hexenal, hexanal, (*E*,*E*)-2,4-hexadienal, and (*E*)-2-heptenal, were markedly reduced in both *ics* mutant lines (Figure 6C).

To further explore the broader metabolic consequences of *ICS* disruption, we performed an untargeted HPLC-MS analysis of semi-polar metabolites from tomato leaves. PCA again showed a clear separation between *ics* mutants and wild type, with PC1 explaining 40.3% of the variance (Figure 6B), confirming that ICS loss alters primary and secondary metabolism beyond the volatilome (Figure 6A).

Semi-polar metabolomic analysis revealed contrasting patterns of metabolite accumulation in the *ics* mutants (Figure 6D). Increased metabolites included DL-N-acetyltryptophan, DL-phenylalanine, D-arginine, DL-citrulline and caffeoylquinic acid. Phenylalanine accumulation may reflect metabolic adjustments associated with impaired SA biosynthesis, although it does not demonstrate compensatory activation of the phenylalanine ammonia-lyase (PAL) pathway. Increased N-acetyltryptophan also suggests alterations in tryptophan metabolism.

In contrast, several phenylpropanoid derivatives, including 3-O-feruloyl-D-quinic acid and 1-O-sinapoyl-beta-D-glucose, showed reduced abundance. The IAA conjugate indole-3-acetyl-beta-1-D-glucoside also decreased, indicating differential regulation of tryptophan-related metabolites. Notably, several sugar-related metabolites, including 6-acetyl-D-glucose, D-(+)-galactose and D-glucopyranose, were markedly reduced. Consistently, targeted sugar quantification revealed lower glucose or fructose levels in both mutants (Figure S6). Together, these findings indicate that ICS disruption alters amino acid, phenylpropanoid, auxin-related and carbohydrate metabolism, revealing extensive metabolic reprogramming under basal conditions.

Finally, phytohormone profiling revealed that the metabolic reprogramming associated with ICS loss extended to hormonal homeostasis. Levels of IAA, cytokinins, and jasmonic acid were significantly reduced in the mutant lines (Figure S7). Given the central roles of auxin and cytokinins in plant growth and development, and of jasmonic acid in stress and defence signalling, these alterations may contribute to both the severe developmental phenotype and the impaired defence responses of the *ics* mutants.

### *ICS* Disruption Impairs Translational Activity in Tomato

Transcriptomic analyses revealed an apparent discrepancy in the *ics* mutants. Gene Ontology enrichment showed upregulation of genes associated with photosynthesis, photoprotection, carotenoid metabolism, photosystem assembly and chloroplast function (Figure 2A). However, physiological and metabolomic analyses revealed reduced carotenoid and chloroplast-associated lipid accumulation (Figure 4B), altered chloroplast morphology (Figure 5), decreased sugar levels (Figure S6) and severe developmental defects (Figure 1A), together with impaired photosynthetic performance (Figure 3A). In contrast, downregulated genes were enriched in processes related to translation, ribosome biogenesis, rRNA processing and protein maturation (Figure 2B). These findings suggested that impaired protein synthesis might contribute to the discrepancy between transcriptional activation and the observed physiological and metabolic defects.

To investigate this possibility, polysome profiling was performed using leaf tissue from wild-type and *ics* plants (Figure 7). Analysis of ribosomal fractions revealed substantial alterations in ribosome distribution in the mutants. In the enlarged region of the polysome profile, *ics* plants displayed a clear accumulation of monosomes (80S ribosomes) accompanied by a reduction in polysome-associated fractions compared with wild-type plants (Figure 7A). Quantification of ribosomal fractions confirmed a significant increase in monosome abundance together with a decrease in the polysome-to-monosome ratio (Figure 7C).

**Figure 7.**
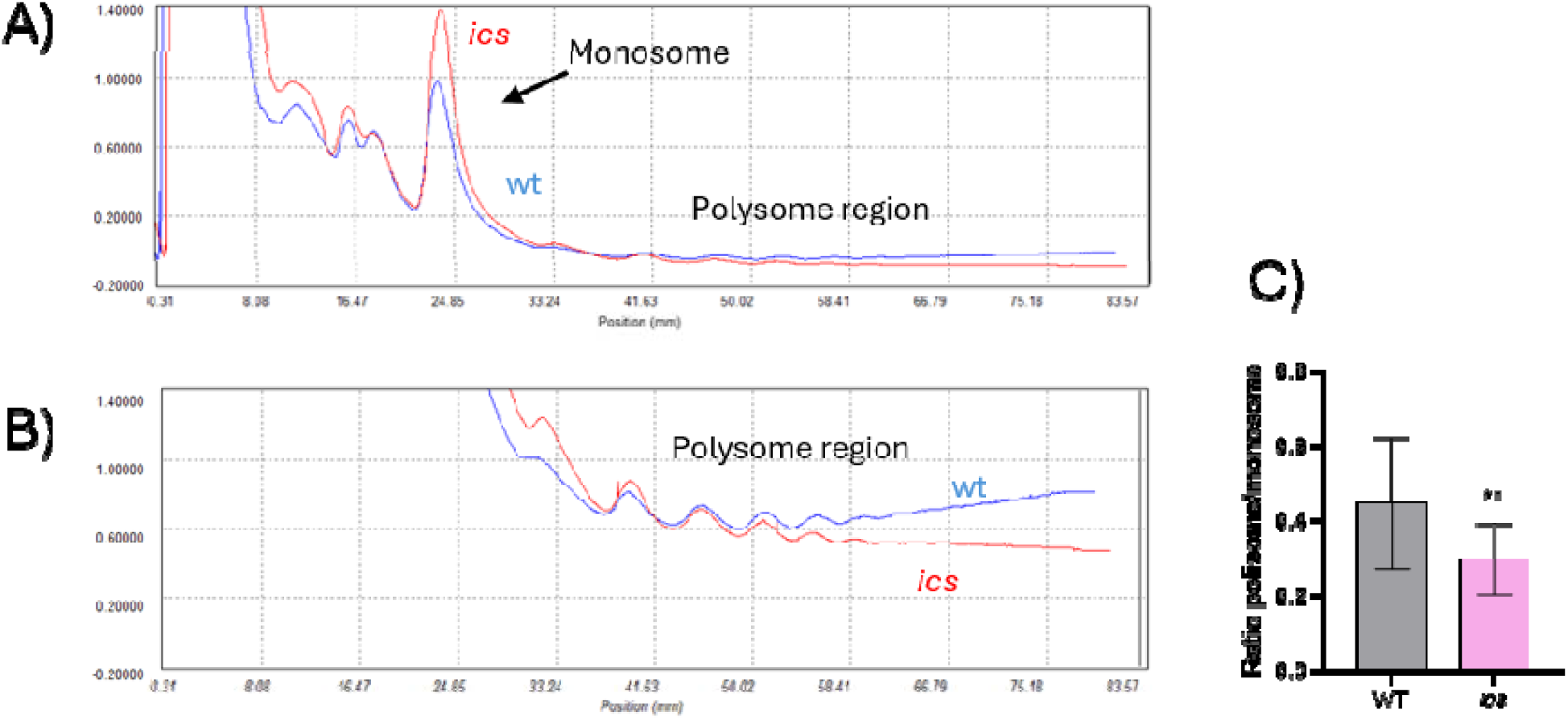
Translational defects in *ics* mutants revealed by polysome profiling. Ribosome profiles were obtained using sucrose density gradient fractionation of leaf extracts from wild-type (WT) and *ics* mutant plants. Absorbance at 254 nm was continuously monitored during fraction collection to visualize ribosomal complexes. **(A)** Magnified view of the monosome (80S) region. **(B)** Magnified view of the polysome region. **(C)** Quantification of the polysome-to-monosome (P/M) ratio. Quantification was performed using four biological replicates from experiments 1 and 2 (n = 4). Data are presented as m an ± SD. Statistical significance was determined by T-test comparison test (\*\**p* < 0.01).

This shift from polysomes toward monosomes is generally considered a well-stablished mark of translation inhibition. Under these conditions, transcripts may still be synthesized and accumulate at high levels, but their efficient translation into proteins is compromised. Consequently, the abundance of specific mRNAs does not necessarily translate into increased protein production.

Taken together, these findings suggest that ICS disruption affects not only salicylic acid biosynthesis and defence signalling but also fundamental processes controlling protein synthesis. The impairment of translation may represent a central mechanism linking the extensive transcriptional reprogramming observed in the mutants with the alterations in chloroplast development, metabolism, and growth.

## Discussion

Isochorismate synthase (ICS) has long been recognized as the major enzyme responsible for pathogen-induced salicylic acid (SA) biosynthesis in *Arabidopsis thaliana* (Katagiri et al., 2002; Garcion et al., 2008). In tomato, SA is also important for defence against bacterial pathogens (Uppalapati et al., 2007; Wang et al., 2025). Previous virus-induced gene silencing (VIGS) experiments showed that ICS silencing reduced pathogen-induced SA accumulation and increased susceptibility to *Pseudomonas syringae* pv. *tomato* (Pst). The bacterial phytotoxin coronatine also suppressed ICS expression and SA-associated defence responses (Uppalapati et al., 2007). However, these studies relied on transient gene silencing and did not examine Pto-mediated effector-triggered immunity (ETI). Thus, the contribution of ICS to ETI and its broader biological functions in tomato remained largely unexplored.

In this study, we generated the first stable CRISPR/Cas9 knockout mutants of tomato *ICS* in a Pto-containing genetic background (Figure 1A), allowing us to investigate its roles in immunity, development and metabolism. Our results demonstrate that *ICS* contributes substantially to pathogen-induced SA accumulation during both virulent and avirulent interactions with Pst, while basal SA levels remain unaffected (Figure 1B). This reduction was particularly pronounced during ETI, when SA accumulation was almost completely abolished and bacterial proliferation significantly increased, indicating that ICS is required for full Pto-mediated resistance (Figure 1D). These findings extend previous observations in tomato based on transient *ICS* silencing by VIGS during virulent Pst infection (Uppalapati et al., 2007) and are consistent with the major contribution of ICS1 to pathogen-induced SA biosynthesis in *Arabidopsis* (Garcion et al., 2008). Importantly, our stable loss-of-function mutants provide evidence that ICS contributes to SA accumulation and full resistance during Pto-mediated ETI in tomato, although its direct contribution to SA biosynthesis remains to be established.

The persistence of residual SA in *ics* mutants suggests that additional biosynthetic routes may contribute to SA production in tomato (Figure 1B). One possible contributor is the phenylalanine-derived pathway, since phenylalanine accumulated in *ics* mutants under basal conditions and during virulent interactions, but not during Pto-mediated ETI (Figure 1C). Although this accumulation does not necessarily indicate increased SA biosynthesis, previous studies have implicated phenylalanine-derived metabolism in tomato defence responses (Umesha et al., 2006; Chang et al., 2008; Zhang et al., 2023). More direct evidence for the contribution of this pathway to SA biosynthesis in tomato was recently provided by Wang et al. (2025), who showed that silencing key enzymes involved in the phenylalanine-derived pathway reduced SA accumulation and *PR1* expression during virulent Pst infection. Together, these observations raise the possibility that this alternative pathway contributes to the residual SA detected in *ics* mutants. During Pto-mediated ETI, however, the SA accumulation was significatively reduced, suggesting that ICS-independent routes do not fully compensate for ICS deficiency under these conditions. Thus, our findings suggest that the contribution of ICS to SA accumulation varies with the type of plant-pathogen interaction, with ICS playing a particularly important role during ETI.

This context-dependent contribution of ICS is consistent with the recent evolutionary diversification of SA biosynthesis across plant species (Ullah et al., 2023; Wang et al., 2025; Zhu et al., 2025; Liu et al., 2026). Recent studies suggest that the canonical ICS-EDS5-PBS3 pathway is evolutionarily restricted to Brassicales (Hong et al., 2025). Moreover, tomato ICS showed limited capacity to promote SA accumulation when co-expressed with Arabidopsis EDS5 and PBS3 in a heterologous system, although this does not exclude its contribution to SA biosynthesis in tomato (Hong et al., 2025). In contrast, a conserved phenylalanine-derived pathway has been identified in seed plants, involving the sequential conversion of benzoyl-CoA into benzyl benzoate, benzyl salicylate and SA (Wang et al., 2025; Zhu et al., 2025; Li & Luo, 2026; Huang & van der Hoorn, 2026). Genetic evidence supporting this pathway in rice and tomato further reinforces its potential contribution to residual SA accumulation in *ics* mutant (Wang et al., 2025; Zhu et al., 2025).

Beyond its effects on immunity, loss of ICS caused a severe developmental phenotype, characterized by strongly reduced growth, environmental sensitivity and failure to complete the life cycle under greenhouse conditions (Figure 1A). Similar growth defects have been reported in the *Arabidopsis thaliana ics1ics2* double mutant, which requires sucrose supplementation for normal maintenance (Garcion et al., 2008). Importantly, exogenous SA application did not restore the developmental phenotype of tomato ics mutants (data not shown), suggesting that ICS deficiency disrupts physiological processes beyond SA-dependent immunity.

A major component of this phenotype appears to involve impaired phylloquinone biosynthesis and photosynthetic function. Isochorismate serves as a precursor not only for SA but also for phylloquinone (vitamin K1), an essential cofactor of Photosystem I that acts as the A1 electron acceptor within the photosynthetic electron transport chain (Reumann, 2013). Consistent with this function, *ics* mutants showed severely reduced phylloquinone levels (Figure 3C) and impaired photosynthetic performance (Figure 3A) despite unchanged chlorophyll content (Figure 3B). Phylloquinone depletion persisted during both virulent and avirulent infections (Figure S4), indicating a constitutive defect that is not restored upon pathogen challenge (Figure S4). These coordinated metabolic alterations indicate that ICS deficiency severely compromises phylloquinone biosynthesis and photosynthetic function. Similar developmental and photosynthetic defects have been reported in *Arabidopsis* mutants impaired in phylloquinone biosynthesis, where disruption of vitamin K1 production affects photosynthetic electron transport, Photosystem I activity, and plant development (Gross et al. 2006). Importantly, supplementation with 1,4-dihydroxy-2-naphthoic acid (NA), a downstream precursor that bypasses the ICS-dependent step, partially restored photosynthetic performance (Figure 3D). A comparable rescue by NA has been reported in rice (Wang et al., 2024), further supporting the role of the isochorismate-phylloquinone pathway in maintaining photosynthetic function. Nevertheless, chloroplast dysfunction may also contribute indirectly to reduced pathogen-induced SA accumulation. Therefore, the relative contributions of direct biosynthetic effects and secondary consequences of impaired photosynthesis remain unresolved.

The involvement of tomato ICS in both SA accumulation and phylloquinone-dependent chloroplast function reflects the functional diversification of ICS across plant species. In *Arabidopsis thaliana*, ICS1 is the predominant source of pathogen-induced SA, but it has also been linked to chloroplast function, affecting thylakoid organization, plastoquinone redox status, and state transitions (Gawroński et al., 2013). In contrast, rice *ICS* plays a predominant role in phylloquinone biosynthesis and photosynthetic function, with only minor effects on SA accumulation (Wang et al., 2024). In cotton, distinct ICS family members preferentially contribute to SA-associated defence or phylloquinone biosynthesis (Guo et al., 2023). By comparison, *Nicotiana benthamiana* ICS displays low catalytic activity and appears to play only a limited role in SA production (Yokoo et al., 2018), whereas in sugarcane, ICS positively regulates both SA accumulation and disease resistance (Zeng et al., 2025). Together, these findings highlight the evolutionary plasticity of ICS function. Tomato is particularly notable because its single ICS gene contributes to both pathogen-associated SA accumulation and phylloquinone biosynthesis, functions that are differentially emphasized or partitioned among paralogs in other species. he severe phylloquinone deficiency in ics mutants was accompanied by reduced accumulation of plastoquinone, carotenoids, prenyl lipids and highly unsaturated fatty acids, important components of thylakoid membranes (Figure 4). These metabolic changes coincided with pronounced defects in chloroplast ultrastructure (Figure 5). Confocal microscopy also revealed ring-like chlorophyll autofluorescence patterns, resembling those described in enlarged chloroplasts, including the division-deficient *arc5-2* mutant (Gao et al., 2003; Knoblauch et al., 2024). Such patterns may arise from peripheral thylakoid distribution surrounding a central stromal region (Knoblauch et al., 2024). Although the underlying organization of the *ics* chloroplasts remains to be established, these similarities suggest that the altered fluorescence patterns may reflect changes in chloroplast geometry and internal membrane organization. Similar links between ICS-dependent phylloquinone biosynthesis, photosynthetic electron transport, and chloroplast activity have been reported in *Arabidopsis thaliana* (Gawroński et al., 2013). Furthermore, reduced diacylglycerols and thylakoid-associated fatty acids provide an additional connection between metabolic and ultrastructural defects. Since chloroplast glycerolipids are essential for thylakoid membrane biogenesis and photosynthetic complex assembly (Kobayashi, 2016), these structural alterations may reflect both phylloquinone deficiency and disrupted membrane lipid composition.

The consequences of impaired chloroplast function extended throughout the metabolome. Reduced soluble sugars and other metabolites are consistent with impaired photosynthetic carbon assimilation (Figures S5 and 6D). Because photosynthesis supplies the energy and carbon skeletons required for primary and secondary metabolism, its impairment may contribute to widespread metabolic reprogramming and growth defects (Ort et al., 2015). Moreover, sugars also function as signalling molecules interacting with immune and hormonal pathways. Thus, altered carbohydrate homeostasis may contribute to the transcriptional, metabolic and defence-related changes observed in the mutants.

Consistent with this metabolic disruption, *ics* mutants showed a marked reduction in terpenoids and green leaf volatiles (GLVs), compounds involved in tomato defence against herbivores and pathogens (Farag and Paré, 2002; López-Gresa et al., 2017; Riedlmeier et al., 2017; Pérez-Pérez et al., 2024) (Figure 6C). Several defence-associated secondary metabolites were also reduced (Figures 6D and S2A–E). These alterations may reflect both impaired SA signalling and reduced carbohydrate availability, highlighting the broader contribution of ICS to metabolic homeostasis (Zhong et al., 2021).

Hormonal homeostasis was also affected, with reduced indole-3-acetic acid (IAA) and cytokinin levels in both mutants (Figure S7A,B). Given their roles in plant growth, chloroplast development and photosynthesis (Gomes and Scortecci, 2021; Vanneste et al., 2025; Emery and Kisiala, 2020; Wu et al., 2021), their depletion may contribute to the developmental defects observed (Vylíčilová et al., 2016). Moreover, the interaction of these hormones with SA-dependent defence pathways (Iglesias et al., 2011; Naseem and Muhammad, 2025) suggests that ICS deficiency may disrupt the coordination between growth and immunity.

An intriguing finding was the discrepancy between transcriptional activation and impaired chloroplast function. RNA-seq analyses revealed strong induction of Gene Ontology categories associated with photosynthesis, photosystem assembly, carotenoid metabolism, photoprotection and chloroplast organization (Figure 2), despite clear physiological and metabolic evidence of impaired chloroplast function. One possible explanation is that this transcriptional activation of these genes represents a compensatory response to ICS disruption. Such a response is consistent with chloroplast-to-nucleus retrograde signalling, whereby changes in plastid metabolic activity, redox status or photosynthetic performance induce extensive transcriptional reprogramming of nuclear genes (Chan et al., 2016; Kleine and Leister, 2016). However, photosynthetic metabolism remained severely compromised, suggesting that transcriptional activation alone is insufficient to restore normal chloroplast functionality. An additional factor potentially contributing to this apparent disconnect is the severe depletion of soluble sugars observed in the *ics* mutants. Carbon availability and cellular energy status are major regulators of translational activity, and reductions in sugar availability are frequently associated with impaired ribosome loading and translational repression (Bailey-Serres and Juntawong, 2012; Cho et al., 2012). Therefore, reduced carbon availability in the mutants may contribute to the altered polysome profiles discussed below.

Polysome profiling revealed an additional alteration associated with ICS deficiency (Figure 7). Compared with wild-type plants, ics mutants accumulated significantly more monosomes and fewer polysomes, consistent with reduced global translational activity. Similar alterations in polysome profiles, accompanied by defective ribosome biogenesis and ribosomal stress, have been reported in tomato plants infected with Citrus exocortis viroid (Cottilli et al., 2019). Although the underlying mechanisms may differ, our findings suggest that ICS disruption also affects the translational machinery. Consequently, reduced translational activity may contribute to the discrepancy between the transcriptional induction of photosynthesis-related genes and the persistent metabolic and physiological defects. However, whether the translation of specific photosynthetic transcripts is directly affected remains to be determined.

Rather than representing a classical growth-defence trade-off, the phenotype of the *ics* mutants illustrates how growth and immunity can become simultaneously compromised when a shared metabolic node is disrupted. Because plant growth and immune responses depend on overlapping carbon, energy and biosynthetic resources (Huot et al., 2014; Karasov et al., 2017), ICS deficiency may compromise growth and defence through combined effects on SA accumulation, chloroplast function and metabolic homeostasis.

Collectively, our findings reveal that the single tomato ICS gene contributes to both pathogen-induced SA accumulation and phylloquinone-dependent chloroplast function. Its disruption causes extensive physiological, metabolic and translational alterations, accompanied by impaired growth and disease resistance. The contribution of ICS to SA accumulation is particularly pronounced during Pto-mediated ETI, although direct biosynthetic effects cannot yet be distinguished from the indirect consequences of chloroplast dysfunction. Together, these results expand our understanding of the species-specific functions of ICS and its contribution to immunity, photosynthesis and cellular homeostasis in tomato.

## Materials and Methods

### Generation of CRISPR/Cas9-mediated *ICS* knockout tomato lines

Tomato (*Solanum lycopersicum* cv. Rio Grande) plants were used throughout this study. CRISPR/Cas9-mediated disruption of ICS (Solyc06g071030.2) was performed using the GoldenBraid 3.0 system (Vázquez-Vilar et al., 2017). Four sgRNAs targeting exons 2 and 3 were designed using Benchling, selected for high predicted on-target efficiency and minimal off-target activity, and assembled with the Cas9 cassette into the pDGB3α1 vector carrying kanamycin resistance and DsRED markers (Figure S1).

The construct was introduced into *Agrobacterium tumefaciens* LBA4404 and used for tomato cotyledon transformation following Ellul et al. (2003). Transgenic plants were selected on kanamycin-containing medium and screened for mutations. Mutant and regenerated WT plants were maintained in vitro under identical conditions on medium supplemented with 20 g L⁻¹ sucrose.

### Bacterial Strain and Inoculation Procedures

Two strains of *Pseudomonas syringae* pv. *tomato* DC3000 (Pst) were used to establish virulent and avirulent interactions with tomato cv. Rio Grande. The Pst DC3000 strain bearing deletions in the *avrPto* and *avrPtoB* genes was used to establish the virulent interaction, whereas Pst DC3000 was used to trigger Pto-mediated effector-triggered immunity (ETI). The bacterial strains were kindly provided by Dr. Selena Giménez (Centro Nacional de Biotecnología, Madrid, Spain) (Lin and Martin, 2005; Ntoukakis et al., 2009). Bacterial cultures and plant inoculations were performed according to established protocols (López-Gresa et al., 2018), with modifications. Briefly, bacterial cells were grown and resuspended to an optical density of 0.1 at 600 nm (OD_600_), generating a stock bacterial suspension. This suspension was subsequently diluted 1:25 in the inoculation solution containing 0.05% Silwet L-77. Four-week-old tomato plants were inoculated by spraying the bacterial suspension onto the aerial parts of the plants. Each Magenta vessel containing four plants was sprayed for 30 s, ensuring uniform coverage of the plant surfaces.

### RNA Extraction and RT-qPCR

The RNA extraction and retro transcription to cDNA of tomato leaves were carried out using column kits based on silica membranes (Macherey-Nagel GmbH, Germany) following the manufacturer’s protocol. cDNA from a microgram of RNA was obtained using a PrimeScript RT reagent kit (Perfect Real Time, Takara Bio Inc., Otsu, Shiga, Japan) following its instructions. qPCRs were performed as previously described (Campos et al. 2014). In each 96-well plate, a reaction took place in a final volume of 10 µL. SYBR R Green PCR Master Mix (Applied Biosystems) was used as the fluorescence marker and actin gene as the endogenous reference gene. Relative gene expression levels were calculated using the ΔΔCt method and are represented in the figures as fold changes relative to the corresponding reference condition indicated in each experiment. The primers used were: PR1 CCGTGCAATTGTGGGTGT/ GAGTTGCGCCAGACTACTTGAGT, P23/PR5 TTC GAG’GTA CGC AAC TG/TGC ATT GAT GAC CCA TGT TT.

### RNAseq

Leaf samples were collected from three individual four-week-old WT, *ics1* and *ics2* plants grown in vitro. RNA sequencing was performed by Macrogen (Seoul, Republic of Korea) using an Illumina NovaSeq 6000 platform (150 bp paired-end reads), generating approximately 60 million reads per sample.

Reads were trimmed using Cutadapt, quality-checked with FastQC and mapped to the tomato reference genome (SL4.0) using STAR. Uniquely mapped reads were quantified using STAR and SAMtools with ITAG4.0 annotation. Differential expression analysis was performed using DESeq2, and functional enrichment was assessed using PANTHER and the gprofiler2 R package (Kolberg et al., 2020).

### Salicylic Acid (SA) Analysis

For the analysis of total SA levels, 0.25 g of frozen, homogenized leaf tissue was extracted in methanol containing 0.25 mM *o*-anisic acid as an internal standard and β-glucosidase was used to hydrolyse SA conjugates. The mixture was sonicated for 10 minutes and centrifuged, and the supernatant was dried under nitrogen flow. SA quantification followed the protocol described by Vázquez Prol et al. (2021), with calibration curves created using authentic standards.

### Jasmonic Acid (JA), Auxin and Citokininis Analysis

Jasmonic acid (JA), auxins and cytokinins were extracted from 100 mg of powdered tomato leaves using 80% methanol containing 1% acetic acid and isotopically labelled internal standards. Extracts were purified using Oasis HLB solid-phase extraction cartridges (Waters), as described by Seo et al. (2011). Phytohormones were separated by UHPLC on an Accucore C18 column and quantified using a Q Exactive Orbitrap mass spectrometer (Thermo Fisher Scientific) operating in targeted selected ion monitoring (tSIM) mode, following Pérez-Pérez et al. (2026). Quantification was performed using calibration curves and the corresponding deuterium-labelled internal standards.

### Analysis of Specialized Metabolites by LC-HESI-MS

For metabolomic profiling, freeze-dried leaf tissue from wild-type (*Rio Grande*) and the two *ics* mutant lines (*ics1* and *ics2*) was finely powdered under liquid nitrogen. Semi-polar metabolites were extracted from 3 mg of lyophilized material using 75% methanol acidified with 0.1% (v/v) formic acid and containing formononetin (0.5 µg mL⁻¹) as an internal standard. Samples were extracted for 30 min at room temperature under continuous shaking (20 Hz) using an MM20 mixer mill (Domel), followed by centrifugation at 20,000 × *g* for 20 min. The resulting supernatants were filtered through 0.2-µm PTFE membrane filter vials (Whatman) prior to chromatographic analysis. Liquid chromatography high-resolution mass spectrometry (LC-HRMS) using an Ultimate HPLC (Dionex) and a Q-Exactive Hybrid Quadrupole-Orbitrap mass spectrometer (Thermo Fisher Scientific, USA), was performed as previously described (Tito A. et al 2026). Data were acquired in both positive and negative ionization modes using a heated electrospray ionization (HESI) source. All solvents were LC-MS grade. Relative metabolite abundances were calculated by normalizing peak areas to the internal standard. Compound identification was based on accurate mass measurements, chromatographic retention times, and MS/MS fragmentation patterns, and was supported by authentic reference standards when available, public spectral repositories, *in-house* databases, and previously published metabolomic datasets. Monoisotopic masses were additionally verified using the PubChem database. Chromatographic data were processed with Xcalibur Quan v4.7 (Thermo Fisher Scientific). Three independent biological replicates were analyzed per genotype, and statistical significance was assessed by one-way ANOVA.

### LC-APCI-HRMS analysis of non-polar compounds

Non-polar metabolites were extracted from 5 mg of lyophilized, homogeneously ground leaf tissues using 2 mL of a 25% (v/v) methanol, 50% (v/v) chloroform, and 25% (v/v) 50 mM Tris-HCl solution, spiked with 15 μg/mL DL-α-tocopherol acetate as the internal standard. LC-HRMS system consisted of a Dionex Ultimate HPLC (Dionex) coupled to a Q-Exactive Hybrid Quadrupole-Orbitrap Mass Spectrometer (Thermo Fisher Scientific). Ionization was obtained by Atmospheric Pressure Chemical Ionization (APCI) source, operating in positive and negative ionization mode. Masses were detected in the 110–1100 m/z range and LC-Ms condition were applied as previously described (Sulli et al 2023). Chlorophylls were quantified in term of μg/g of dry weight (DW), by their on-line absorption PDA (photo Diode Array) spectra, as described in (Fantini et al. 2019).

### Comparative untargeted metabolomics

Untargeted metabolomics was carried out using raw LC-HRMS files collected from both semi-polar and non-polar fractions, processed with Compound Discoverer v3.3 (Thermo Fisher Scientific), applying the Untargeted Metabolomics Workflow with statistics, using three replicates each sample, and setting parameters as previously reported (Akhgari A. et al., 2026). Metabolic features were filtered based on peak quality and reproducibility, aligning chromatograms using the ChromAlign algorithm (Sadygov et al 2006), with the RT between 0.8 and 15 or 28 min for LC-APCI-MS and LC-HESI-MS analysis, respectively. Peaks showing peak rating ≥ 6 in at least three samples, group CV [%]) ≤ 28 in at least one sample group, and group area ≥3 × 10^6^, were selected. MS2 annotated compounds were assigned by the exact mass (Dppm ≤ 5), using MS2 fragments match with mzCloud (confidence ≥ 60/100 between the experimental and reference spectra) or *in silico* fragmentation using FISh (Fragment Ion Search) scoring (≥ 40). Differentially abundant (DA) tentatively identified compounds were further selected by p-value ≤ 0.05 within WT and both *ics* lines (Akhgari A. et al 2026). The list of tentatively identified compounds with the detailed information is reported in Table S1 and S2, for non-polar and semi-polar metabolites, respectively.

### Volatile Organic Compound (VOC) Analysis

Volatile organic compounds (VOCs) were extracted and analysed following a modified protocol based on López-Gresa et al. (2017). Briefly, 100 mg of frozen powdered tomato leaf tissue was transferred to 10-mL glass vials containing 1 mL of 6 M CaCl₂ and 100 μL of 750 mM EDTA (pH 7.5). Samples were sealed immediately and sonicated for 5 min before VOC extraction by headspace solid-phase microextraction (HS-SPME). Volatile compounds were subsequently analysed by GC-MS as described in Balanzá et al., 2025, and chromatograms were processed using Agilent Enhanced ChemStation software, which enabled compound identification through comparison of mass spectra and retention times with the integrated reference library. Quantification was performed using Agilent-developed targeted methods based on characteristic quantifier ions and retention times.

### Phylloquinone Quantification

Phylloquinone (vitamin K₁) was extracted from 100 mg of fresh tomato leaf tissue following a modified protocol of Verberne et al. (2007). Samples were homogenized in 1 mL of isopropanol containing 3.25 µM menaquinone K₂ as an internal standard, heated at 80 °C for 5 min and extracted with isopropanol:hexane (1:2, v/v). Following sonication and incubation at 80 °C for 10 min, water was added and the hexane phase was collected after centrifugation and evaporated under nitrogen.

Dried extracts were resuspended in ethanol and treated with SnCl₂ (60 mg mL⁻¹ in ethanol:HCl, 1:1, v/v) for 10 min. Phylloquinone was quantified by HPLC-fluorescence (excitation 240 nm, emission 430 nm) using methanol:water (95:5, v/v) as the mobile phase. Quantification was based on calibration curves with authentic standards, using menaquinone K₂ to control analytical variability.

### Polysome profiling

Polysome profiling was performed following Mustroph et al. (2009), with modifications. Leaf tissue (200 mg) was homogenized in 0.4 mL of extraction buffer containing 200 mM Tris-HCl (pH 9.0), 200 mM KCl, 5 mM EGTA, 35 mM MgCl₂, 4% Triton X-100, 1% sodium deoxycholate, 5 mM DTT and 0.5 mg mL⁻¹ heparin. Extracts (300 µL) were loaded onto 10-50% sucrose gradients and ultracentrifuged at 273,000 × g for 1 h 40 min at 4 °C. Gradients were fractionated using a BioComp Gradient Station, and ribosomal profiles were continuously monitored by UV absorbance at 260 nm.

### Confocal microscopy

Confocal imaging was performed using a Leica Stellaris WLL laser scanning confocal microscope equipped with a HC PL APO CS2 40×/1.25 glycerol immersion objective. Samples were excited using the 488 nm laser line, and fluorescence emission was collected between 650 and 700 nm. All images were acquired using identical microscope settings for all samples to allow direct comparison.

### LIQOR600 fluorescence measurements

Photosystem II (PSII) performance was assessed in intact leaves by measuring chlorophyll fluorescence using a LI-6800 Portable Photosynthesis System (LI-COR Biosciences, Lincoln, NE, USA). Measurements were performed on attached, fully expanded leaves to evaluate photosynthetic performance.

### Treatment with 1,4-dihydroxy-2-naphthoic acid in *in vitro* conditions

To evaluate the effect of phylloquinone precursor supplementation, plants were grown on Murashige and Skoog (MS) medium supplemented with 0.1 mM 1,4-dihydroxy-2-naphthoic acid (NA), prepared as a stock solution in dimethyl sulfoxide (DMSO). The final concentration of DMSO in the culture medium was 0.1% (v/v). Control plants were grown on MS medium supplemented with 0.1% (v/v) DMSO only.

Plants were cultivated under standard growth conditions for one month. Photosystem II performance was subsequently assessed by measuring chlorophyll fluorescence parameters using a LI-6800 Portable Photosynthesis System (LI-COR Biosciences, Lincoln, NE, USA).

### Transmission electronic microscopy for chloroplast visualization

Selected leaf fragments were fixed overnight in Karnovsky’s solution (2.5% glutaraldehyde and 2% paraformaldehyde in 0.1 M phosphate buffer). A vacuum system was used to facilitate the fixative penetration into the tissues. Following this, the tissues were post-fixed in 2% osmium tetroxide. The samples were then rinsed, dehydrated through a graded ethanol series, and embedded in epoxy resin.

To ensure precise visualization of the internal structures, specimens were properly oriented during embedding. Ultrathin sections (0.06-0.08 µm) were cut with an Ultracut UC-6 (Leica microsystems, Wetzlar, Germany) with a diamond knife, stained with lead citrate (Reynold’s solution) and examined under a transmission electron microscope FEI Tecnai G2 Spirit BioTwin (Thermo Fisher Scientific company, Oregon, USA). All images were acquired with a Xarosa digital camera (EMSIS GmbH, Münster, Germany) controlled by Radius software (Version 2.1).

## Acknowledgements

We acknowledge the Plant Hormone Quantification Service at the IBMCP (Valencia, Spain) for performing the hormone analyses. We also acknowledge Javier Forment for performing the RNA-seq analysis and Mario Soriano from Instiute Príncipe Felipe for providing the TEM microscopy images. We also want to thank Marisol Gascón Irún for her help and support with confocal microscopy images.

This work was supported by grants PID2020-116765RB-I00 and PID2023-152361OB-I00 funded by MCIN/AEI/10.13039/501100011033, and grants PROMETEU/2021/056 and CIPROM/2024/41 from Generalitat Valenciana. Additional funding was provided by the CIDEGENT programme of Generalitat Valenciana (CIDEXG/2022/27, awarded to B.B.-P.) and the Spanish Ministry of Science, Innovation and Universities (PID2022-142412NB-I00, awarded to B.B.-P.). This work also received support from institutional grant CEX2025-001626-S (Severo Ochoa Centre of Excellence), funded by MCIU/AEI/10.13039/501100011033. J.P.-P. was supported by an EMBO Scientific Exchange Grant (No. 11432) and a predoctoral fellowship from the Spanish Ministerio de Universidades (FPU21/00259).

## Author contributions

**J.P.-P.**: Methodology, Investigation, Validation, Formal analysis, Data curation, Visualization, Writing - original draft, Writing - review & editing. **M.S.**: Methodology, Investigation, Formal analysis, Data curation, Writing - review & editing. **B.B.**: Investigation, Writing - review & editing. **M.R.-R.**: Methodology, Investigation, Writing - review & editing. **F.V.-S.**: Methodology, Formal analysis, Data curation, Visualization, Writing - review & editing. **G.D.**: Methodology, Formal analysis, Data curation, Supervision, Writing - review & editing. **I.R.**: Formal analysis, Data curation, Visualization, Writing - review & editing. **M.P.L.-G.**: Conceptualization, Supervision, Project administration, Funding acquisition, Writing - review & editing. **P.L.**: Conceptualization, Supervision, Project administration, Funding acquisition, Writing - review & editing.

## Data availability

All other data supporting the findings of this study are available within the article and its Supporting Information.

## References

1. Akhgari, A., Nava, M., Leitner, R., Demurtas, O. C., Twyman, R. M., Becue, I., Van Poucke, C., Van Droogenbroeck, B., Brillmann, M., Oksman-Caldentey, K.-M., Diretto, G., and Rischer, H. (2026). Industrial-scale production of basil hairy roots in a 1000-L bioreactor and advanced downstream processing for the preparation of cosmetic ingredients. New Biotechnology, 95, 103–112. 10.1016/j.nbt.2026.08.005

2. Bailey-Serres, J., and Juntawong, P. (2012). Dynamic light regulation of translation status in Arabidopsis thaliana. Frontiers in Plant Science 3, 66. 10.3389/fpls.2012.00066

3. Balanzá, M., Vázquez-Prol, F., Rodrigo, I., Bellés, J.M., Vera-Sirera, F., López-Gresa, M.P., and Lisón, P. (2025). Salicylic acid modulates volatile organic compound profiles during CEVd infection in tomato plants. Metabolites 15, 102. 10.3390/metabo15020102

4. Cameron, R. K., Paiva, N. L., Lamb, C. J., and Dixon, R. A. (1999). Accumulation of salicylic acid and PR-1 gene transcripts in relation to the systemic acquired resistance (SAR) response induced by *Pseudomonas syringae* pv. *tomato* in *Arabidopsis*. Physiological and Molecular Plant Pathology, 55(2), 121–130. 10.1006/pmpp.1999.0214

5. Campos, L., Granell, P., Tárraga, S., López-Gresa, M. P., Conejero, V., Bellés, J. M., Rodrigo, I., and Lisón, P. (2014). Salicylic acid and gentisic acid induce RNA silencing-related genes and plant resistance to RNA pathogens. Plant Physiology and Biochemistry, 77, 35–43. 10.1016/j.plaphy.2014.01.016

6. Chan, K.X., Phua, S.Y., Crisp, P., McQuinn, R., and Pogson, B.J. (2016). Learning the languages of the chloroplast: retrograde signalling and beyond. Annual Review of Plant Biology 67, 25–53. 10.1146/annurev-arplant-043015-112231

7. Chang, A., Lim, M.H., Lee, S.W., Robb, E.J., and Nazar, R.N. (2008). Tomato phenylalanine ammonia-lyase gene family, highly redundant but strongly underutilized. Journal of Biological Chemistry 283, 33591–33601. 10.1074/jbc.M804428200

8. Chen, H., Pan, B., Zhang, S., Li, X., Zhang, Y., Gao, K., Chen, D., Wang, L., Jiang, T., Luo, C., and Huang, C. (2025). Recent advances in biosynthesis and bioactivity of plant caffeoylquinic acids. Current Issues in Molecular Biology, 47(11), 942. 10.3390/cimb47110942

9. Cho, Y.H., Hong, J.W., Kim, E.C., and Yoo, S.D. (2012). Regulatory functions of SnRK1 in stress-responsive gene expression and in plant growth and development1Plant Physiology 158(4):1955-64, doi: 10.1104/pp.111.189829.

10. Cottilli, P., Belda-Palazón, B., Adkar-Purushothama, C.R., Perreault, J.P., Schleiff, E., Rodrigo, I., Ferrando, A., and Lisón, P. (2019). Citrus exocortis viroid causes ribosomal stress in tomato plants. Nucleic Acids Research 47, 8649–8661. 10.1093/nar/gkz679.

11. El-Azaz J, Cánovas FM, Barcelona B, Ávila C, de la Torre F. Deregulation of phenylalanine biosynthesis evolved with the emergence of vascular plants. Plant Physiol. 2022;188(1):134–150. doi:10.1093/plphys/kiab454

12. Emery, R.J.N., and Kisiala, A. (2020). The roles of cytokinins in plants and their response to environmental stimuli. Plants 9, 1158. 10.3390/plants9091158.

13. Fantini, E., Sulli, M., Zhang, L., Aprea, G., Jiménez-Gómez, J. M., Bendahmane, A., Perrotta, G., Giuliano, G., and Facella, P. (2019). Pivotal roles of cryptochromes 1a and 2 in tomato development and physiology. Plant Physiology, 179(2), 732–748. 10.1104/pp.18.00793

14. Farag MA, Paré PW. (2002)C6-Green leaf volatiles trigger local and systemic VOC emissions in tomato. Phytochemistry.61(5):545–554. 10.1016/S0031-9422(02)00240-6.

15. Gao, H., Kadirjan-Kalbach, D., Froehlich, J. E., and Osteryoung, K. W. (2003). ARC5, a cytosolic dynamin-like protein from Arabidopsis, is required for chloroplast division and mediates squeeze-off of plastid envelopes. Proceedings of the National Academy of Sciences, 100(7), 4328–4333. 10.1073/pnas.0530206100

16. Gao, Q. M., Zhu, S., Kachroo, P., and Kachroo, A. (2015). Signal regulators of systemic acquired resistance. Frontiers in Plant Science, 6, 228. 10.3389/fpls.2015.00228

17. Garcion, C., Lohmann, A., Lamodière, E., Catinot, J., Buchala, A., Doermann, P., & Métraux, J. P. (2008). Characterization and biological function of the ISOCHORISMATE SYNTHASE2 gene of Arabidopsis. Plant physiology, 147(3), 1279–1287. 10.1104/pp.108.119420

18. Gawroński, P., Górecka, M., Bederska, M., Rusaczonek, A., Ślesak, I., Kruk, J., and Karpiński, S. (2013). Isochorismate synthase 1 is required for thylakoid organization, optimal plastoquinone redox status, and state transitions in *Arabidopsis thaliana*. Journal of Experimental Botany 64, 3669–3679. 10.1093/jxb/ert203

19. Gomes, G.L.B., and Scortecci, K.C. (2021). Auxin and its role in plant development: Structure, signalling, regulation and response mechanisms. Plant Biology 23, 894–904. 10.1111/plb.13303.

20. Gross, J., Cho, W.K., Lezhneva, L., Falk, J., Krupinska, K., Shinozaki, K., Seki, M., Herrmann, R.G., and Meurer, J. (2006). A plant locus essential for phylloquinone (vitamin K1) biosynthesis originated from a fusion of four eubacterial genes. Plant Journal 47, 295–306. 10.1111/j.1365-313X.2006.02790.x

21. Guo, Z., Zhang, W., Ruan, C., Kong, J., Wang, F., Wang, G., and Guo, W. (2023). Two cotton isochorismate synthase genes are biased functionally in regulating salicylic acid and phylloquinone biosynthesis. Environmental and Experimental Botany 216, 105540. 10.1016/j.envexpbot.2023.105540

22. Havaux, M. (1998). Carotenoids as membrane stabilizers in chloroplasts. Trends in Plant Science, 3(4), 147–151. 10.1016/S1360-1385(98)01200-X

23. Hong, K., Nakano, M., Tang, Y., Jeanguenin, L., Kang, W., Wang, Y., Zuo, L., Li, P., He, J., Jiang, W., Huang, R., Matsui, H., Wang, Y., Nakagami, H., Li, B., Li, X., Xie, K., Fukushima, K., Guo, L., … Tsuda, K. (2025). Emergence of isochorismate-based salicylic acid biosynthesis within Brassicales. Proceedings of the National Academy of Sciences, 122(29), e2506170122. 10.1073/pnas.2506170122

24. Huang, J., and van der Hoorn, R. A. L. (2026). The ancestral salicylic acid biosynthesis pathway in plants. Trends in Plant Science, 31(1), 17–19. 10.1016/j.tplants.2025.08.004

25. Huot, B., Yao, J., Montgomery, B.L., and He, S.Y. (2014). Growth–defence tradeoffs in plants: a balancing act to optimize fitness. Molecular Plant 7, 1267–1287. 10.1093/mp/ssu049

26. Iglesias, M.J., Terrile, M.C., and Casalongué, C.A. (2011). Auxin and salicylic acid signallings counteract the regulation of adaptive responses to stress. Plant Signalling & Behavior 6, 452–454. 10.4161/psb.6.3.14676

27. Jones, J.D.G. & Dangl, J.L. (2006). The plant immune system. Nature 444, 323–329. 10.1038/nature05286.

28. Karasov, T.L., Chae, E., Herman, J.J., and Bergelson, J. (2017). Mechanisms to mitigate the trade-off between growth and defence. Plant Cell 29, 666–680. 10.1105/tpc.16.00931

29. Katagiri, F., Thilmony, R., and He, S.Y. (2002). The *Arabidopsis thaliana*–*Pseudomonas syringae* interaction. The Arabidopsis Book 1, e0039. 10.1199/tab.0039

30. Kim, H.U., van Oostende, C., Basset, G.J.C., and Browse, J. (2008). The AAE14 gene encodes the Arabidopsis o-succinylbenzoyl-CoA ligase that is essential for phylloquinone synthesis and photosystem-I function. The Plant Journal 54, 272–283. 10.1111/j.1365-313X.2008.03416.x

31. Kleine, T., and Leister, D. (2016). Retrograde signalling: organelles go networking. Biochimica et Biophysica Acta 1857, 1313–1325. 10.1016/j.bbabio.2016.03.017

32. Klessig, D.F., Choi, H.W. & Dempsey, D.A. (2018). Systemic Acquired Resistance and Salicylic Acid: Past, Present, and Future. Mol. Plant-Microbe Interact. 31, 871–888. 10.1094/MPMI-03-18-0067-CR

33. Knoblauch, J., Waadt, R., Cousins, A. B., and Kunz, H.-H. (2024). Probing the in situ volumes of Arabidopsis leaf plastids using three-dimensional confocal and scanning electron microscopy. The Plant Journal, 117(2), 332–341. 10.1111/tpj.16554

34. Kolberg, L., Raudvere, U., Kuzmin, I., Vilo, J., and Peterson, H. (2020). gprofiler2 – an R package for gene list functional enrichment analysis and namespace conversion toolset g:Profiler. F1000Research 9, 709. 10.12688/f1000research.24956.2

35. Li, Y., and Luo, J. (2026). Unraveling the central puzzle in salicylic acid biosynthesis from phenylalanine. Journal of Integrative Plant Biology, 68(4), 849–851. 10.1111/jipb.70016

36. Lin, N. C., and Martin, G. B. (2005). An avrPto/avrPtoB mutant of Pseudomonas syringae pv. tomato DC3000 does not elicit Pto-mediated resistance and is less virulent on tomato. Molecular Plant-Microbe Interactions, 18(1), 43–51. 10.1094/MPMI-18-0043

37. Liu, Y., Wu, M., Li, X., and Zhang, Y. (2026). Conserved and divergent: Salicylic acid biosynthesis and signalling pathways across the plant kingdom. Molecular Plant 19, 587–605. 10.1016/j.molp.2026.02.007

38. Liu, Y., Xu, L., Wu, M., Wang, J., Qiu, D., Lan, J., Lu, J., Zhang, Y., Li, X., … Zhang, Y. (2025). Three-step biosynthesis of salicylic acid from benzoyl-CoA in plants. Nature, 645, 201–207. 10.1038/s41586-025-09185-7

39. Loake, G., and Grant, M. (2007). Salicylic acid in plant defence—the players and protagonists. Current Opinion in Plant Biology, 10(5), 466–472. 10.1016/j.pbi.2007.08.008

40. Lonjon, F., Lai, Y., Askari, N., Aiyar, N., Bundalovic-Torma, C., Laflamme, B., Wang, P. W., Desveaux, D., and Guttman, D. S. (2024). The effector-triggered immunity landscape of tomato against *Pseudomonas syringae*. Nature Communications, 15(1), 5102. 10.1038/s41467-024-49425-4

41. López-Gresa, M. P., Lisón, P., Campos, L., Rodrigo, I., Rambla, J. L., Granell, A., Conejero, V., and Bellés, J. M. (2017). A non-targeted metabolomics approach unravels the VOCs associated with the tomato immune response against *Pseudomonas syringae*. Frontiers in Plant Science, 8, 1188. 10.3389/fpls.2017.01188

42. López-Gresa, M. P., Payá, C., Ozáez, M., Rodrigo, I., Conejero, V., Klee, H., Bellés, J. M., & Lisón, P. (2018). A new role for green leaf volatile esters in tomato stomatal defence against *Pseudomonas syringae* pv. tomato. Frontiers in Plant Science, 9, Artículo 1855. 10.3389/fpls.2018.01855

43. Mustroph, A., Zanetti, M.E., Jang, C.J., Holtan, H.E., Repetti, P.P., Galbraith, D.W., Girke, T., and Bailey-Serres, J. (2009). Profiling translatomes of discrete cell populations resolves altered cellular priorities during hypoxia in *Arabidopsis*. Proceedings of the National Academy of Sciences of the United States of America 106, 18843–18848. 10.1073/pnas.0906131106

44. Naseem, M., and Muhammad, K. (2025). Genome editing approaches to harness cytokinin– salicylic acid crosstalk for plant protection. Frontiers in Genome Editing 7, 1687599. 10.3389/fgeed.2025.1687599

45. Ngou, B. P. M., Ahn, H.-K., Ding, P., and Jones, J. D. G. (2021). Mutual potentiation of plant immunity by cell-surface and intracellular receptors. Nature, 592(7852), 110–115. 10.1038/s41586-021-03315-7

46. Nićiforović, N., and Abramovič, H. (2014). Sinapic acid and its derivatives: Natural sources and bioactivity. Comprehensive Reviews in Food Science and Food Safety, 13(1), 34–51. 10.1111/1541-4337.12041

47. Ntoukakis, V., Mucyn, T. S., Gimenez-Ibanez, S., Chapman, H. C., Gutierrez, J. R., Balmuth, A. L., Jones, A. M., & Rathjen, J. P. (2009). Host inhibition of a bacterial virulence effector triggers immunity to infection. Science, 324(5928), 784–787. 10.1126/science.1169430

48. Ort, D.R., Merchant, S.S., Alric, J., Barkan, A., Blankenship, R.E., Bock, R., Croce, R., Hanson, M.R., Hibberd, J.M., Long, S.P., Moore, T.A., Moroney, J., Niyogi, K.K., Parry, M.A.J., Peralta-Yahya, P.P., Prince, R.C., Redding, K.E., Spalding, M.H., van Wijk, K.J., Vermaas, W.F.J., von Caemmerer, S., Weber, A.P.M., Yeates, T.O., Yuan, J.S., and Zhu, X.G. (2015). Redesigning photosynthesis to sustainably meet global food and bioenergy demand. Proceedings of the National Academy of Sciences USA 112, 8529–8536. 10.1073/pnas.1424031112

49. Pérez-Pérez, J., Minguillón, S., Kabbas-Piñango, E., Payá, C., Campos, L., Rodríguez-Concepción, M., Espinosa-Ruiz, A., Rodrigo, I., Bellés, J. M., López-Gresa, M. P., and Lisón, P. (2024). Metabolic crosstalk between hydroxylated monoterpenes and salicylic acid in tomato defence response against bacteria. Plant Physiology, 195(3), 2323–2338. 10.1093/plphys/kiae148

50. Pérez-Pérez, J., Brito-Gutiérrez, P., Santiago, A., Sanmartín, M., Matus, J. T., Sulli, M., Diretto, G., Vera-Sirera, F., Rodrigo, I., López-Gresa, M. P., and Lisón, P. (2026). Hydroxylated monoterpenes activate tomato defence pathways against bacteria to trigger jasmonate-dependent self-amplifying immunity. Plant Stress 24, 101594. 10.1016/j.stress.2026.101594

51. Rekhter, D., Lüdke, D., Ding, Y., Feussner, K., Zienkiewicz, K., Lipka, V., Wiermer, M., Zhang, Y., and Feussner, I. (2019). Isochorismate-derived biosynthesis of the plant stress hormone salicylic acid. Science, 365(6452), 498–502. 10.1126/science.aaw1720

52. Reumann, S. (2013). Biosynthesis of vitamin K1 (phylloquinone) by plant peroxisomes and its integration into signalling molecule synthesis pathways. In Subcellular Biochemistry, Vol. 69, pp. 213–229. 10.1007/978-94-007-6889-5_12

53. Riedlmeier, M., Ghirardo, A., Wenig, M., Knappe, C., Koch, K., Georgii, E., Dey, S., Parker, J. E., Schnitzler, J. P., & Vlot, A. C. (2017). Monoterpenes Support Systemic Acquired Resistance within and between Plants. The Plant cell, 29(6), 1440–1459. 10.1105/tpc.16.00898

54. Romer, J., Gutbrod, K., Schuppener, A., Melzer, M., Müller-Schüssele, S. J., Meyer, A. J., and Dörmann, P. (2024). Tocopherol and phylloquinone biosynthesis in chloroplasts requires the phytol kinase VITAMIN E PATHWAY GENE5 (VTE5) and the farnesol kinase (FOLK). The Plant Cell, 36(4), 1140–1158. 10.1093/plcell/koad316

55. Sadygov, R. G., Maroto, F. M., and Hühmer, A. F. R. (2006). ChromAlign: A two-step algorithmic procedure for time alignment of three-dimensional LC–MS chromatographic surfaces. Analytical Chemistry, 78(24), 8207–8217. 10.1021/ac060923y

56. Shimada, H., Ohno, R., Shibata, M., Ikegami, I., Onai, K., Ohto, M., and Takamiya, K. (2005). Inactivation and deficiency of core proteins of photosystems I and II caused by genetical phylloquinone and plastoquinone deficiency but retained lamellar structure in a T-DNA mutant of Arabidopsis. The Plant Journal 41, 627–637. 10.1111/j.1365-313X.2004.02326.x

57. Soll, J. (1987). α-Tocopherol and plastoquinone synthesis in chloroplast membranes. Methods in Enzymology, 148, 383–392. 10.1016/0076-6879(87)48037-3

58. Spoel, S. H., and Dong, X. (2024). Salicylic acid in plant immunity and beyond. The Plant Cell, 36(5), 1451–1464. 10.1093/plcell/koad329

59. Sulli, M., Dall’Osto, L., Ferrante, P., Guardini, Z., Gomez, R. L., Mini, P., Demurtas, O. C., Aprea, G., Nicolia, A., Bassi, R., & Giuliano, G. (2023). Generation and physiological characterization of genome-edited Nicotiana benthamiana plants containing zeaxanthin as the only leaf xanthophyll. Planta, 258(5), 93. 10.1007/s00425-023-04248-3

60. Tito, A., Vardaro, E., De Lucia, A., Falanga, D., De Musis, C., Colantuono, A., Cefariello, A., Ricci, L., Vitiello, G., Arra, I., Di Lorenzo, R., Diretto, G., Demurtas, O. C., Sulli, M., Pappas, A., Greco, G., Laneri, S., and Colucci, M. G. (2026). *Hyssopus officinalis* L. (Lamiaceae) Cell Culture Extract Modulates Epidermal Lipid-Related and Differentiation Markers. Cells, 15(14), 1300. 10.3390/cells15141300

61. Torrens-Spence, M. P., Bobokalonova, A., Carballo, V., Glinkerman, C. M., Pluskal, T., Shen, A., and Weng, J.-K. (2019). PBS3 and EPS1 complete salicylic acid biosynthesis from isochorismate in Arabidopsis. Molecular Plant, 12(12), 1577–1586. 10.1016/j.molp.2019.11.005

62. Ullah, C., Chen, Y.-H., Ortega, M. A., and Tsai, C.-J. (2023). The diversity of salicylic acid biosynthesis and defence signalling in plants: Knowledge gaps and future opportunities. Current Opinion in Plant Biology, 72, 102349. 10.1016/j.pbi.2023.102349

63. Umesha, S. (2006). Phenylalanine ammonia lyase activity in tomato seedlings and its relationship to bacterial canker disease resistance. Phytoparasitica 34, 68–71. 10.1007/BF02981341

64. Uppalapati, S. R., Ishiga, Y., Wangdi, T., Kunkel, B. N., Anand, A., Mysore, K. S., and Bender, C. L. (2007). The phytotoxin coronatine contributes to pathogen fitness and is required for suppression of salicylic acid accumulation in tomato inoculated with *Pseudomonas syringae* pv. *tomato* DC3000. Molecular Plant-Microbe Interactions, 20(8), 955–965. 10.1094/MPMI-20-8-0955

65. Vanneste, S., Pei, Y., and Friml, J. (2025). Mechanisms of auxin action in plant growth and development. Nature Reviews Molecular Cell Biology 26, 648–666. 10.1038/s41580-025-00851-2.

66. Vázquez-Vilar, M., Quijano-Rubio, A., Fernández-del-Carmen, A., Sarrion-Perdigones, A., Ochoa-Fernandez, R., Ziarsolo, P., Blanca, J., Granell, A., and Orzaez, D. (2017). GB3.0: A platform for plant bio-design that connects functional DNA elements with associated biological data. Nucleic Acids Research 45(4), 2196–2209. 10.1093/nar/gkw1326

67. Verberne, M. C., Sansuk, K., Bol, J. F., Linthorst, H. J. M., and Verpoorte, R. (2007). Vitamin K1 accumulation in tobacco plants overexpressing bacterial genes involved in the biosynthesis of salicylic acid. Journal of Biotechnology 128, 72–79. 10.1016/j.jbiotec.2006.09.005.

68. Vylíčilová, H., Husičková, A., Spíchal, L., Srovnal, J., Doležal, K., Plíhal, O., et al. (2016). C2-substituted aromatic cytokinin sugar conjugates delay the onset of senescence by maintaining the activity of the photosynthetic apparatus. Phytochemistry 122, 22–33. 10.1016/j.phytochem.2015.12.001

69. Wang, L., Li, Q., Zhang, A., Zhou, W., Jiang, R., Yang, Z., Yang, H., Qin, X., Ding, S., Lu, Q., Wen, X., and Lu, C. (2017). The phytol phosphorylation pathway is essential for the biosynthesis of phylloquinone, which is required for photosystem I stability in Arabidopsis. Molecular Plant 10, 183–196. 10.1016/j.molp.2016.12.006

70. Wang, Y., Jin, G., Song, S., Jin, Y., Wang, X., Yang, S., Shen, X., Gan, Y., Wang, Y., Li, R., Liu, J.-X., Hu, J., and Pan, R. (2024). A peroxisomal cinnamate:CoA ligase-dependent phytohormone metabolic cascade in submerged rice germination. Developmental Cell, 59(11), 1363–1378.e4. 10.1016/j.devcel.2024.03.023

71. Wang, Z., Yang, G., Zhang, D., Li, G., Qiu, J. L., and Wu, J. (2024). Isochorismate synthase is required for phylloquinone, but not salicylic acid biosynthesis in rice. aBIOTECH, 5(4), 488–496. 10.1007/s42994-024-00166-4

72. Wang Y, Song S, Zhang W, Deng Q, Feng Y, Tao M, et al. 2025. Deciphering phenylalanine-derived salicylic acid biosynthesis in plants. Nature, 645(8079), 208–217. 10.1038/s41586-025-09280-9

73. Wildermuth, M.C., Dewdney, J., Wu, G., and Ausubel, F.M. (2001). Isochorismate synthase is required to synthesize salicylic acid for plant defence. Nature 414, 562–565. 10.1038/35107108.

74. Willis, D. K., and Kinscherf, T. G. (2009). Population dynamics of *Pseudomonas syringae* pv. *tomato* strains on tomato cultivars Rio Grande and Rio Grande-Pto under field conditions. Journal of Phytopathology, 157(4), 219–227. 10.1111/j.1439-0434.2008.01481.x

75. Wu, W., Du, K., Kang, X., et al. (2021). The diverse roles of cytokinins in regulating leaf development. Horticulture Research 8, 118. 10.1038/s41438-021-00558-3.

76. Yang, J., Shen, G., & Wu, J. (2025). Jasmonic acid and salicylic acid transcriptionally regulate CuRe1 in cultivated tomato to activate resistance to parasitization by dodder *Cuscuta australis*. Plant Diversity, 47(3), 511–521. 10.1016/j.pld.2025.03.003

77. Yang, W., Xu, X., Li, Y., Wang, Y., Li, M., Wang, Y., Ding, X., and Chu, Z. (2016). Rutin-mediated priming of plant resistance to three bacterial pathogens initiating the early SA signal pathway. PLoS ONE, 11(1), e0146910. 10.1371/journal.pone.0146910

78. Yokoo, S., Inoue, S., Suzuki, N., Amakawa, N., Matsui, H., Nakagami, H., Takahashi, A., Arai, R., and Katou, S. (2018). Comparative analysis of plant isochorismate synthases reveals structural mechanisms underlying their distinct biochemical properties. Bioscience Reports, 38(2), BSR20171457. 10.1042/BSR20171457

79. Yu, L., Shen, W., Fan, J., Sah, S. K., Mavraganis, I., Wang, L., Gao, P., Gao, J., Zheng, Q., Meesapyodsuk, D., Yang, H., Li, Q., Zou, J., and Xu, C. (2023). A chloroplast diacylglycerol lipase modulates glycerolipid pathway balance in *Arabidopsis*. The Plant Journal, 115(2), 335–350. 10.1111/tpj.16228

80. Yuan, M., Jiang, Z., Bi, G., Nomura, K., Liu, M., Wang, Y., Cai, B., Zhou, J.-M., He, S. Y., and Xin, X.-F. (2021). Pattern-recognition receptors are required for NLR-mediated plant immunity. Nature, 592(7852), 105–109. 10.1038/s41586-021-03316-6

81. Zeng, Z., Zhu, X., Chen, J., Zhang, S., Zhou, S., Li, K., and Shen, W. (2025). Genomic and functional analysis of isochorismate synthase genes in sugarcane and their roles in disease resistance. The Crop Journal, 13(4), 1210–1223. 10.1016/j.cj.2025.05.013

82. Zhang, F., Wang, J., Li, X., Zhang, J., Liu, Y., Chen, Y., Yu, Q., and Li, N. (2023). Genome-wide identification and expression analyses of phenylalanine ammonia-lyase gene family members from tomato (*Solanum lycopersicum*) reveal their role in root-knot nematode infection. Frontiers in Plant Science 14, 1204990. 10.3389/fpls.2023.1204990

83. Zhong, Q., Hu, H., Fan, B., Zhu, C., and Chen, Z. (2021). Biosynthesis and roles of salicylic acid in balancing stress response and growth in plants. International Journal of Molecular Sciences, 22(21), 11672. 10.3390/ijms222111672

84. Zhu, B., Zhang, Y., Gao, R., Wu, Z., Zhang, W., Zhang, C., Zhang, P., Ye, C., Yao, L., Jin, Y., Mao, H., Tou, P., Huang, P., Zhao, J., Zhao, Q., Liu, C.-J., and Zhang, K. (2025). Complete biosynthesis of salicylic acid from phenylalanine in plants. Nature, 645(8079), 218–227. 10.1038/s41586-025-09175-9

