## Supplemental information for "The Tomato Isochorismate Synthase Coordinates both Salicylic Acid-Mediated Immunity and Phylloquinone-Dependent Chloroplast Function"

ATGGCTGTAGGTGTAAGGCACTGCACCATAAGGTTCATGGAACTTGAATCATCATCACTAATGAAATGTTTGCTCTTGTCTTCACCTCTATATAGAAAGCAATCAGT
TACCGACATCCACATTCCGTGACGTGGTATTCCAAGTACCTTGAACTTAGTAGTAGTGATTACTTTACAAACGAGAACAGAAGTGGAGATATATCTTTCGTTAGTCA

EXON 1

10 20 30 40 50 60 70 80 90 100

TCACTTCTCCAACTCTTCTACTCAAAGATATAATCAATGTTGTTCATTGTCGATGAATGGATGCCAAGGCGATTCAAGAGCTCCTATTGGAATCGTTGAAACTCGAA
AGTGAAGAGGTTGAGAAGATGAGTTTCTATATTAGTTACAACAAGTAACAGCTACTTACCTACGGTTCCGCTAAGTTCTCGAGGATAACCTTAGCAACTTTGAGCTT

icsl mutation
guide 3
ics2 mutation
EXON 1
EXON 2

110 120 130 140 150 160 170 180 190 200 210

CTTTGCCTGCAGTTTCTAGCCCTGCATTGGCCATGGAACGTCTCAATTCTGCCATCTCCGATATGATTAAGTCCGATCCTCCGCCTTACGATTCCGGCATTATTCGC
GAAACGGACGTCAAAGATCGGGACGTAACCGGTACCTTGCAGAGTTAAGACGGTAGAGGCTATACTAATTCAGGCTAGGAGGCGGAATGCTAAGGCCGTAATAAGCG

icsl mutation
guide 2
guide 1
ics2 mutation
EXON 2

220 230 240 250 260 270 280 290 300 310 320

CTTGAGGTGCCAATTGAAGAGCAGATTGAATCACTTGAGTGGCTTCATGCCCAGAACCATCTCCTCCTCCCTCGCTGCTTCTTCTCCGGTCGAAGAGTTGCTTCTGA
GAACTCCACGGTTAACTTCTCGTCTAACTTAGTGAACTCACCGAAGTACGGGTCTTGGTAGAGGAGGAGGGAGCGACGAAGAAGAGGCCAGCTTCTCAACGAAGACT

guide 4
EXON 3

330 340 350 360 370 380 390 400 410 420

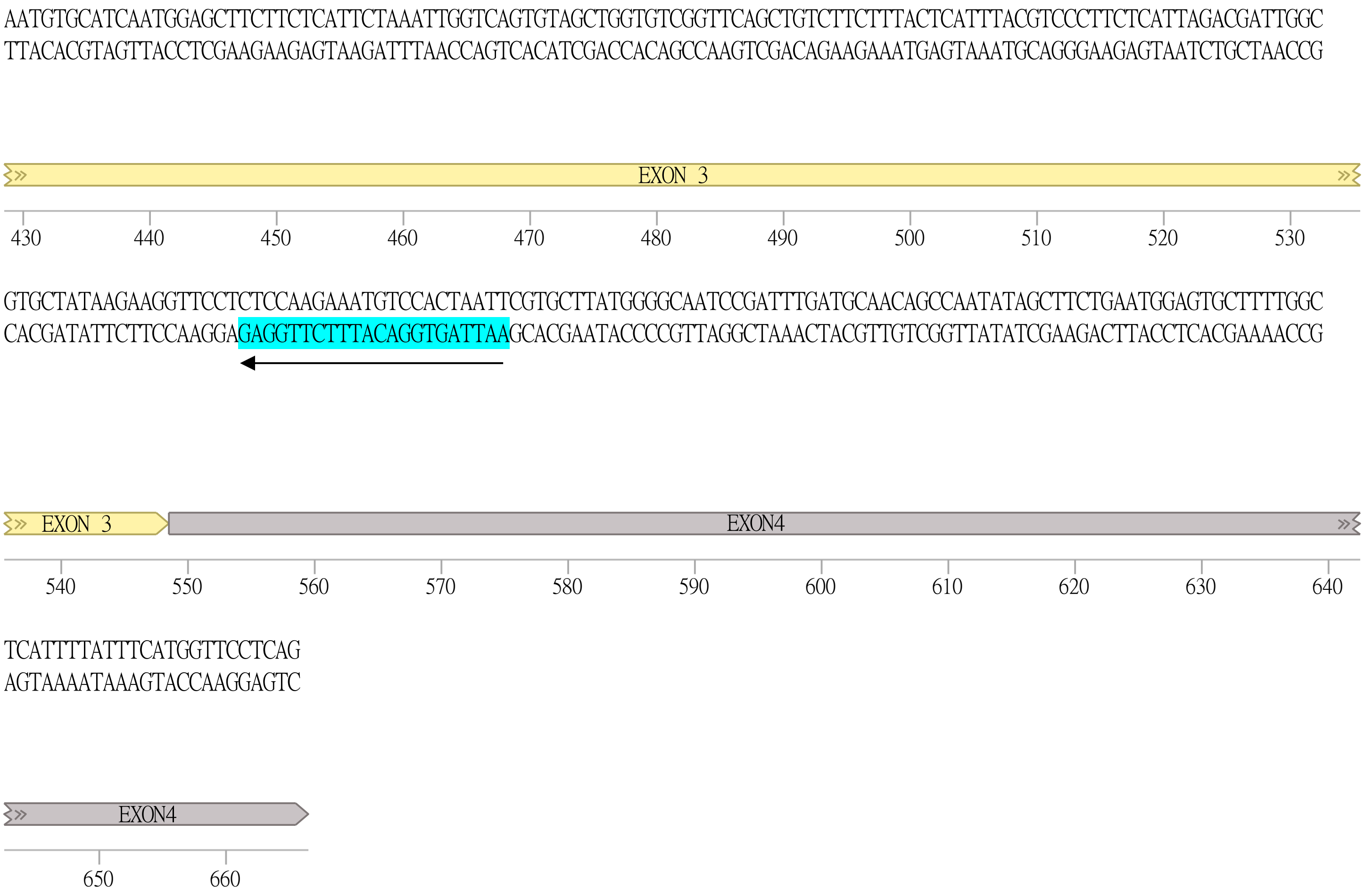


**Figure S1. Schematic representation of the amplified cDNA regions encompassing the *ICS* CRISPR target sites.** The figure shows the nucleotide sequences of the amplified genomic regions, together with their corresponding complementary strands and the genomic position of each fragment. Exons 1–4 are represented as coloured arrows along the genomic sequence, with nucleotide positions indicated below each region. The positions of the four CRISPR guide RNAs (guides 1–4) are marked with coloured arrows, while the binding sites of the primers used for genotyping are indicated by light blue coloured boxes. The regions containing the mutations identified in *ics1* and *ics2* are highlighted in orange and grey, respectively, within exon 2 and 3.

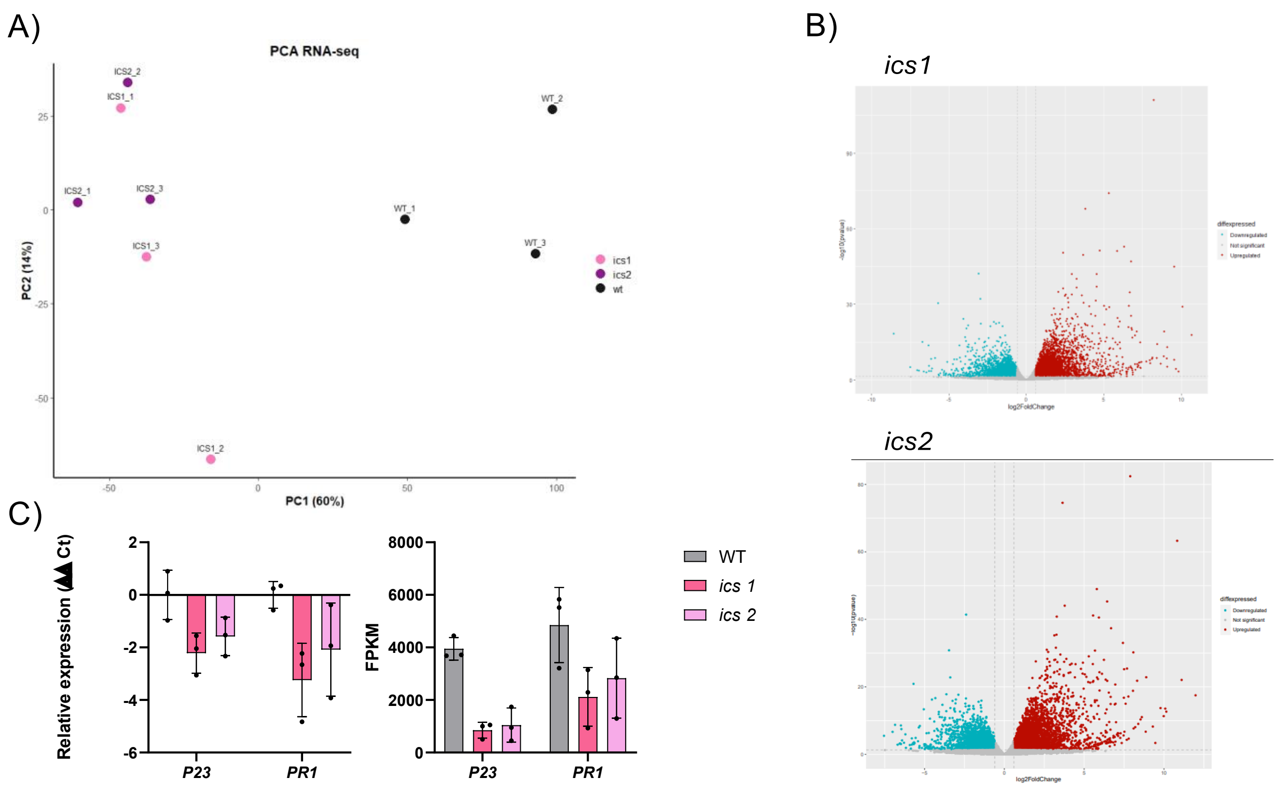


**Figure S2. Transcriptomic reprogramming in CRISPR-generated *ics* mutant tomato plants under basal conditions. (A)** Principal component analysis (PCA) of RNA-seq data showing a clear separation between wild-type (WT), *ics1* and *ics2* **(B)** Volcano plots comparing differentially expressed genes (DEGs) between WT and *ics1* (top right) and WT and *ics2* (bottom right), highlighting significantly upregulated (red) and downregulated (blue) genes. **(C)** Quantification of selected representative genes PR1 and P23 associated with defensive responses in FPKM and relative expression.

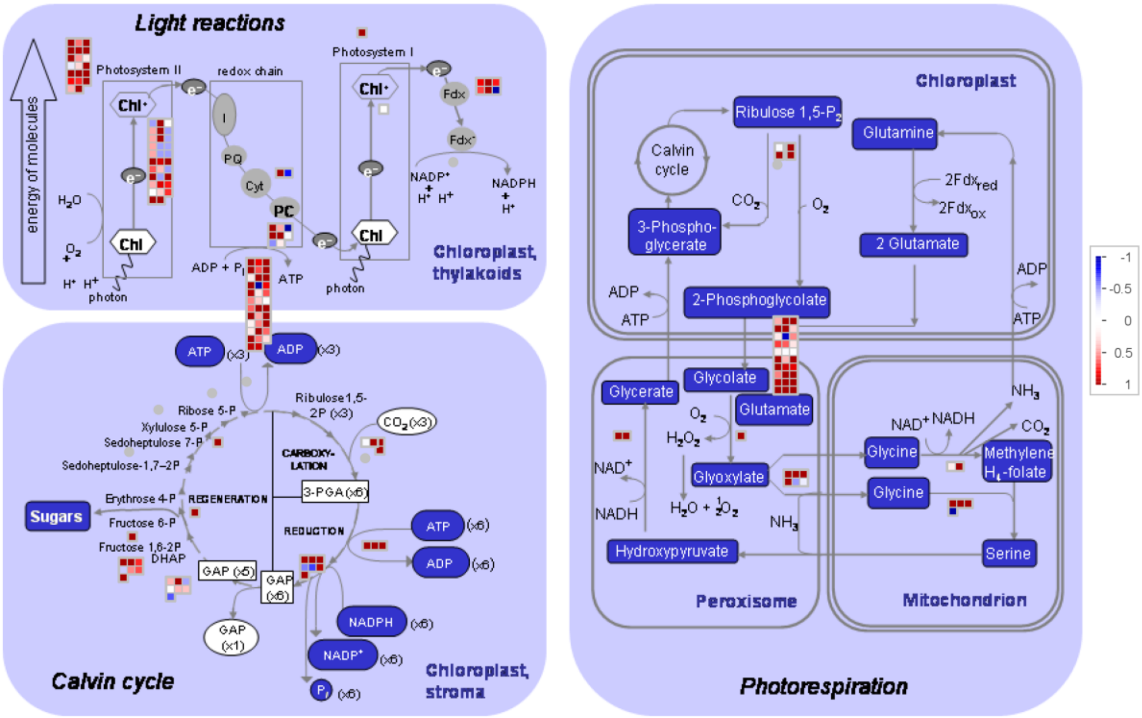


**Figure S3. MapMan visualization of changes in genes associated to primary carbon metabolism, photosynthesis, and photorespiration in *ics* mutants compared with wild-type plants**. Differentially expressed genes are displayed within the corresponding metabolic pathways, including the light reactions, Calvin cycle, and photorespiration. Metabolites are represented as colored squares, where blue indicates increased accumulation (induction) and red indicates decreased accumulation (repression) relative to the wild type. Color intensity is proportional to the magnitude of the change, as indicated by the scale.

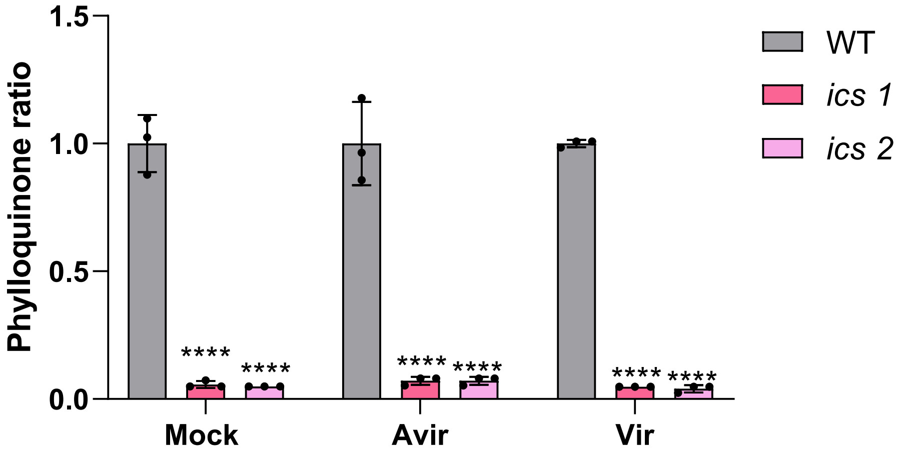


**Figure S4. Levels of phylloquinone in *ics* mutants under virulent and avirulent infections.** Data are presented as mean ± standard deviation (SD) of 3 biological replicates. Data are expressed relative to the WT mean, normalized to 1, to illustrate changes in metabolite levels in *ics* mutant plants. Statistical significance was determined relative to WT using one-way ANOVA followed by Tukey's multiple comparison test (****$p$< 0.001).

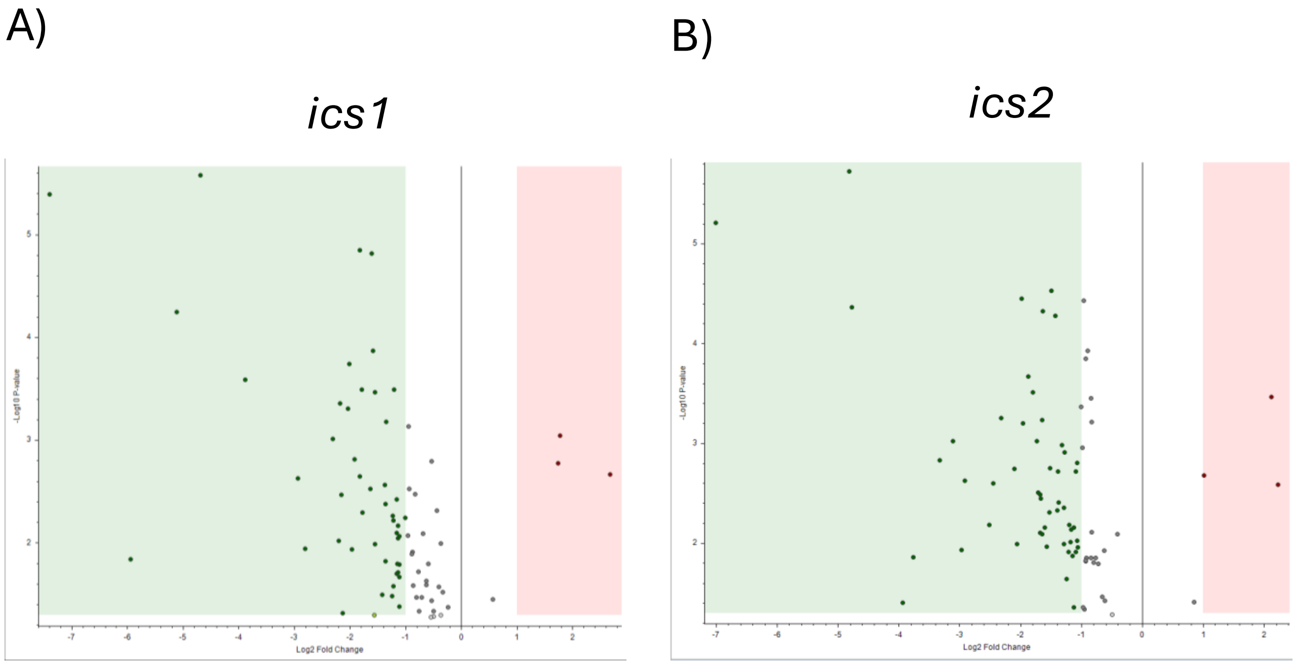


**Figure S5. Comparative untargeted metabolomic analysis of non-polar compounds in *ics* and wt leaves. (A)** Distribution of the DA non-polar metabolites showing significantly altered levels within WT and ics1 **(A)** and ics2 **(B)** lines. Red or green dots represent, respectively, metabolites accumulated at lower and upper levels in ics plants compared to the WT ($p < 0.05$, log2 FC D/P > 1).

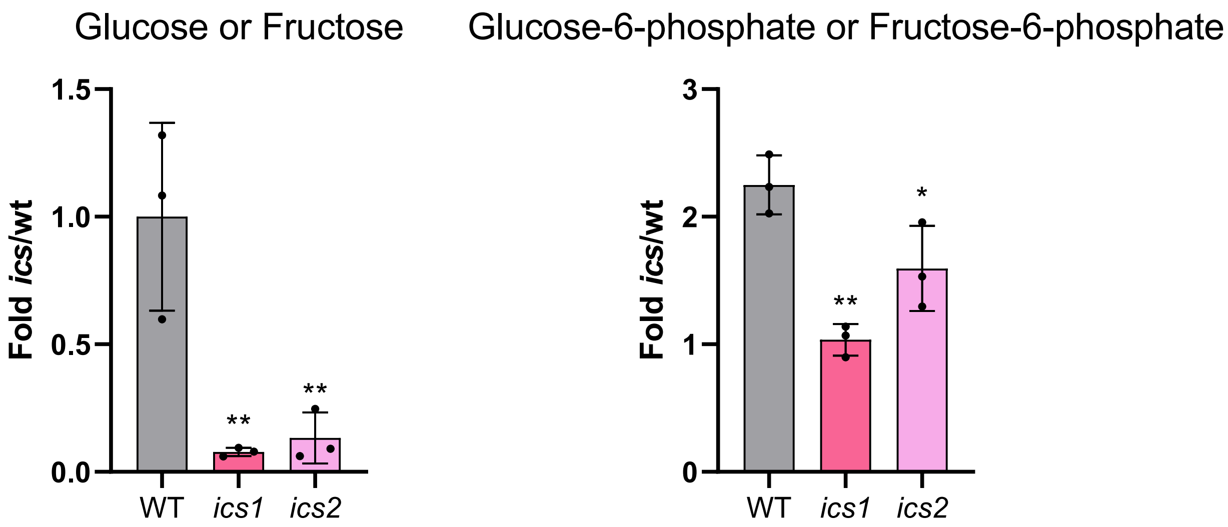


**Figure S6. Levels of sugars in *ics* mutants.** Levels of glucose (or fructose) and glucose-6-phosphate (or fructose-6-phosphate) in wild-type (WT) and *ics* mutant tomato plants (*ics1* and *ics2*). Data are expressed relative to the WT mean, normalized to 1, to illustrate changes in metabolite levels in *ics* mutant plants. Bars represent the mean (n=3) ± SE. Asterisks denote significant differences relative to WT ($p$ < 0.01).

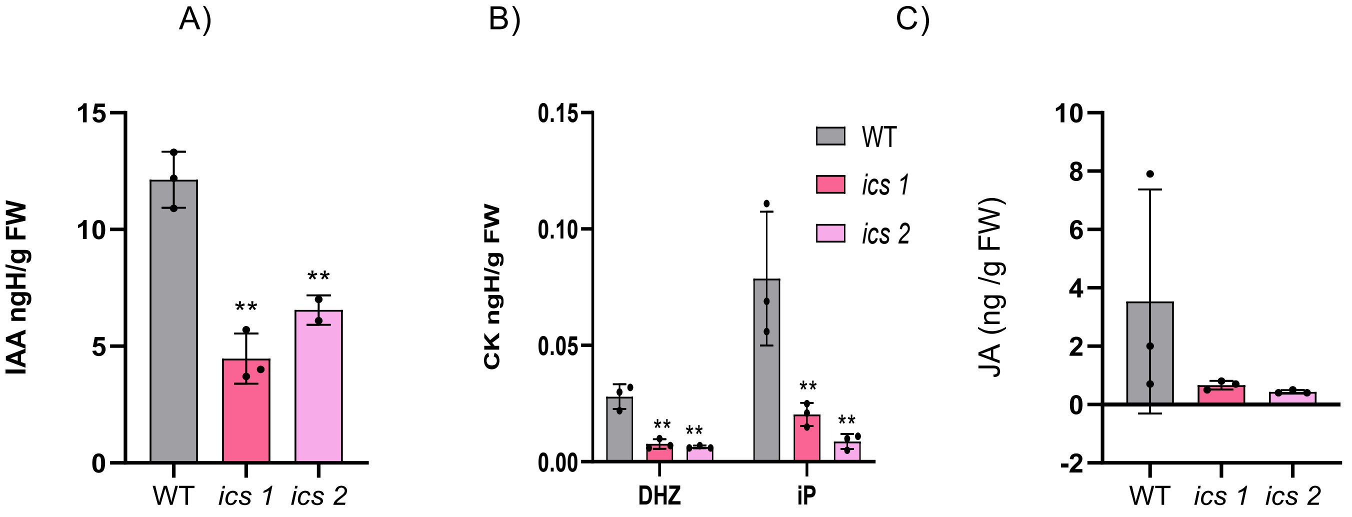


**Figure S7. Quantification of phytohormones in *ics* mutans plants. (A-B)** Targeted hormonal profiling of auxin (Indole-3-Acetic Acid, IAA) and cytokinins (Dihydrozeatin, DHZ, and $N^6$-Isopentenyladenine, iP) **(C)** Targeted jasmonic acid quantification (ng/g FW) in WT plants and *ics* mutants. Data represent mean ± SD. Statistical significance was determined relative to WT using one-way ANOVA followed by Dunnett's multiple comparison test (*$p < 0.05$, **$p < 0.01$).
